# Single cell plasticity and population coding stability in auditory thalamus upon associative learning

**DOI:** 10.1101/2020.04.06.026401

**Authors:** James Alexander Taylor, Masashi Hasegawa, Chloé Maëlle Benoit, Joana Amorim Freire, Marine Theodore, Dan Alin Ganea, Tingjia Lu, Jan Gründemann

**Affiliations:** Department of Biomedicine, University of Basel, Klingelbergstrasse 50-70, 4056 Basel, Switzerland; Friedrich Miescher Institute for Biomedical Research, Maulbeerstrasse 66, 4058 Basel, Switzerland

**Keywords:** Associative learning, fear conditioning, deep brain imaging, auditory thalamus, medial geniculate body, population coding, multisensory integration

## Abstract

Cortical and limbic brain areas are regarded as centres for learning. However, how thalamic sensory relays participate in plasticity upon associative learning, yet support stable long-term sensory coding remains unknown. Using a miniature microscope imaging approach, we monitor the activity of populations of auditory thalamus (MGB) neurons in freely moving mice upon fear conditioning. We find that single cells exhibit mixed selectivity and heterogeneous plasticity patterns to auditory and aversive stimuli upon learning, which is conserved in amygdala-projecting MGB neurons. In contrast to individual cells, population level encoding of auditory stimuli remained stable across days. Our data identifies MGB as a site for complex neuronal plasticity in fear learning upstream of the amygdala that is in an ideal position to drive plasticity in cortical and limbic brain areas. These findings suggest that MGB’s role goes beyond a sole relay function by balancing experience-dependent, diverse single cell plasticity with consistent ensemble level representations of the sensory environment to support stable auditory perception with minimal affective bias.

## Introduction

Associative learning depends on the reliable integration of sensory stimuli from the environment with specific aversive or appetitive outcomes to shape future behaviours. Many cortical and limbic brain areas have been identified as centres for associative learning. However, how thalamic sensory relays like the auditory thalamus (medial geniculate body, MGB), which provide direct sensory input to these areas, participate in plasticity upon associative learning, yet ensure stable long-term sensory coding remains unknown. Auditory fear conditioning, a well-studied classical conditioning paradigm, identified the amygdala as a core brain area for associative learning of stimulus-predicted (conditioned stimulus, CS, e.g. tone) aversive outcomes (unconditioned stimulus, US, e.g. mild foot shock) (Fanselow and LeDoux, 1999; Maren and Fanselow, 1996; Rogan et al., 1997; Schafe et al., 2000). At its input site, amygdala response plasticity is thought to be driven by synaptic potentiation in basolateral amygdala (BLA) neurons (Humeau et al., 2003; McKernan and Shinnick-Gallagher, 1997). However, early work demonstrated that the higher order MGB, a major auditory input hub to the amygdala (Rogan and LeDoux, 1995), is a site of CS-US integration and plasticity upon fear learning (Apergis-Schoute et al., 2005; Bordi and LeDoux, 1994a, 1994b; Edeline and Weinberger, 1991b, 1991a; Han et al., 2008; Hennevin et al., 1993; Ryugo and Weinberger, 1978). Enhanced responses to conditioned stimuli and increased synaptic drive from presynaptic MGB neurons to the BLA might act as an additional plasticity mechanism for associative fear learning. Nevertheless, the role of MGB in neuronal response plasticity upon fear learning has been controversially discussed (Fanselow and LeDoux, 1999; Maren et al., 2001; Weinberger, 2011) and recent physiological studies of fear conditioning mostly excluded this site of sensory integration and response potentiation upstream of the amygdala and auditory cortex. It is currently unknown if individual MGB neurons exhibit complex response dynamics upon adaptive associative and defensive behaviours and how this potential heterogeneity is balanced with reliable representations of sensory inputs from the environment. Furthermore, we are currently lacking a concept of the ensemble level activity and dynamics in this widely projecting thalamic auditory relay site, which is crucial to delineate the distributed population code underlying associative learning and adaptive defensive behaviours (Headley et al., 2019; Mobbs et al., 2020).

Here we used a combination of deep brain Ca^2+^ imaging, miniature microscopy and fear conditioning in freely moving mice to reveal the response dynamics and plasticity of large populations of auditory thalamus neurons (Ghosh et al., 2011; Grewe et al., 2017; Gründemann et al., 2019). We find that individual auditory thalamus neurons exhibit mixed selectivity of CS and US responses with highly diverse plasticity patterns during associative learning, while the ensemble representation of auditory stimuli remains stable along learning. These findings suggest that auditory thalamus plays a role beyond a classic relay function by balancing experience-dependent plasticity with stable ensemble level representations of the sensory environment to support stable auditory perception with minimal affective bias.

## Results

### Deep brain imaging of auditory thalamus during fear conditioning

We established a gradient-index lens deep brain miniature microscope imaging approach of identified auditory thalamus neuronal populations in freely behaving mice (**Figures 1A and 1B**) (Ghosh et al., 2011; Gründemann et al., 2019). Using genetically-encoded Ca^2+^ sensors (**Figure 1C**, AAV2/5.CaMKII.GCaMP6f) (Chen et al., 2013), we tracked large populations of individual MGB neurons across a four-day auditory fear conditioning paradigm in freely moving mice. Similar to previous reports (Hackett et al., 2016), we found GABAergic fibres, typically originating in inferior colliculus and the thalamic reticular complex (Rouiller et al., 1985; Winer et al., 1996), but virtually no GABAergic somata in MGB (**Figure S1**, see also Jager et al., 2019), indicating that we mainly imaged Ca^2+^ activity of thalamic relay neurons. We were able to follow 97 ± 5 GCaMP6f-expressing neurons per mouse (**Figures 1D and 1E**, N = 19 mice) stably within and across sessions. MGB neurons exhibited diverse, spontaneous activity patterns in freely moving animals (**Figure 1F**) as well as cell-specific responses to pure tone auditory stimuli (**Figure 1G**).

**Figure 1:**
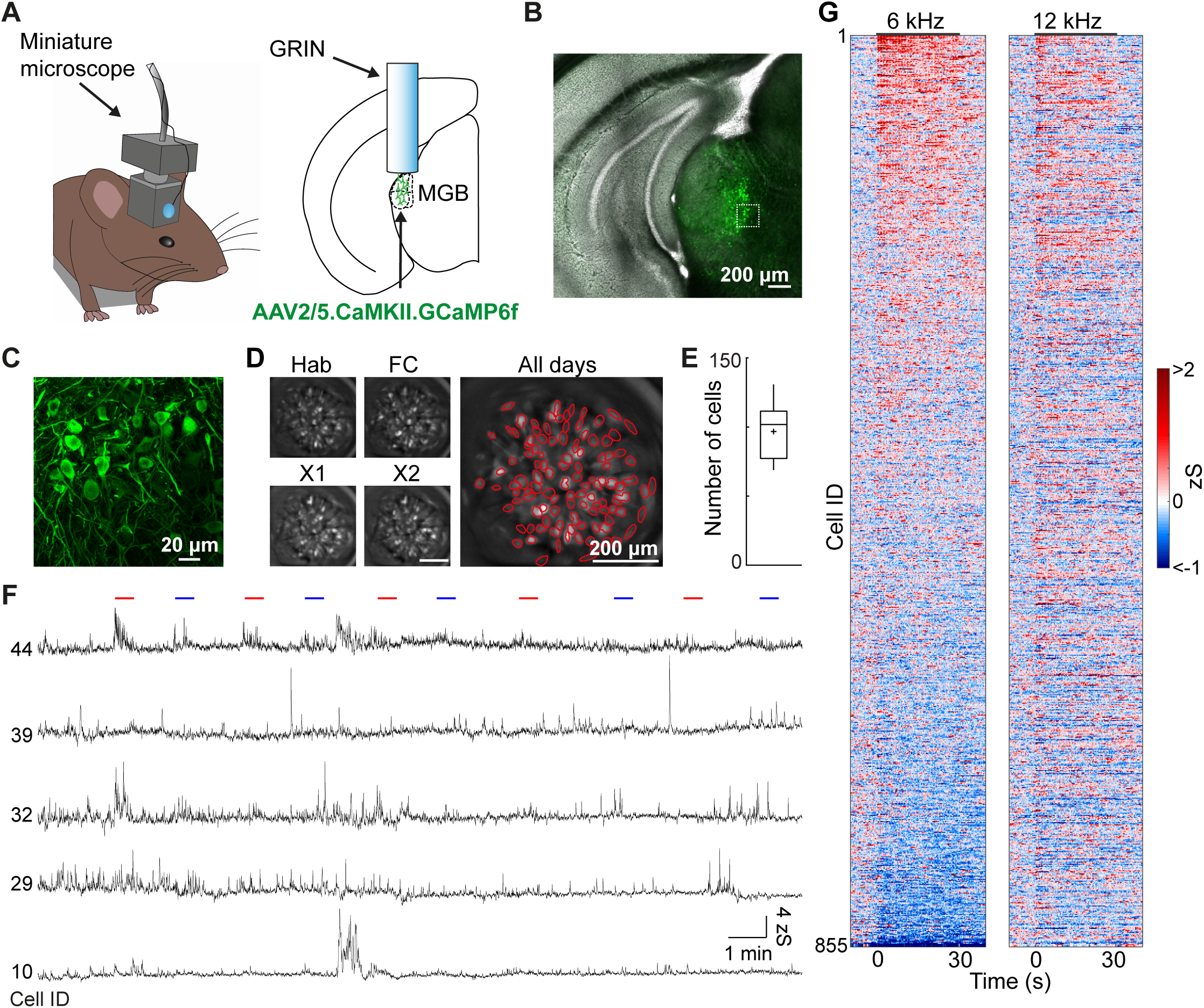
Imaging neuronal activity of auditory thalamus in freely moving mice. **A)** Mouse with a head-mounted miniaturized microscope (left). Location of gradient refractive index (GRIN) lens in the medial geniculate body (MGB, right). **B)** GCaMP6f expression in MGB. **C)** High magnification of GCaMP6f-expressing MGB neurons from B. **D)** Individual motion corrected field of views (maximum intensity projection) of one example animal across a four-day fear conditioning paradigm (Hab, FC, Ext.1, Ext.2) as well as the maximum intensity projection across all days. Red circles indicate selected individual components. **E)** Average number of individual ICs / animal (97 ± 5 neurons, N = 19 mice). **F)** Changes in Ca^2+^ fluorescence of five individual neurons during the habituation session. Lines indicate CS tone presentations (red: 12 kHz, blue: 6 kHz). **G**) Tone responses on habituation day 1 of all recorded MGB neurons in fear conditioning experiments (n = 855 neurons, N = 9 mice).

Next, we used a classical four-day fear conditioning and fear extinction paradigm (**Figures 2A and 2B**) (Herry et al., 2008), in which mice learn to associate a mild foot-shock unconditioned stimulus (US) with a predictive conditioned stimulus (CS+, 6 or 12 kHz pure tones, 200 ms pips, 27 pips per CS). After fear conditioning, mice exhibited enhanced freezing to the CS+ (61% ± 6%, neutral CS-freezing: 42% ± 8%, N = 15, **Figures 2B and S2A**), which extinguished upon repetitive CS+ presentation (Friedman test, p < 0.001, followed by Dunn’s multiple comparisons test, Extinction 1 early vs. Extinction 1 late p < 0.01, Extinction 1 early vs. Extinction 2 late p < 0.001, Extinction 2 early vs. Extinction 2 late p < 0.05). During fear conditioning, the total neuronal population (**Figure 2C**) as well as individual MGB neurons (**Figures 2D and 2E**) were strongly responsive to both the CS+ and the US. The proportion of US responsive neurons (75% ± 5%) was significantly higher than the proportion of CS+ (27% ± 3%) and CS-neurons (20% ± 2%, N = 9 mice, see Methods for classification of responsive neurons), while similar proportions of neurons were responsive to the CS+ and the CS- (**Figures 2F, S2C, and S2D**, Friedman test, p < 0.001 followed by Dunn’s multiple comparisons test CS+ vs. CS-p > 0.05, CS+ vs. US p < 0.05, CS- vs. US p < 0.001). Furthermore, we found mixed selectivity in subpopulations of neurons that were responsive to combinations of tones and foot shocks, yet they were not enriched beyond chance level in the total population (**Figure 2G**). These multisensory neurons were spatially intermingled in MGB and not locally clustered (**Figures 2H-J**).

**Figure 2:**
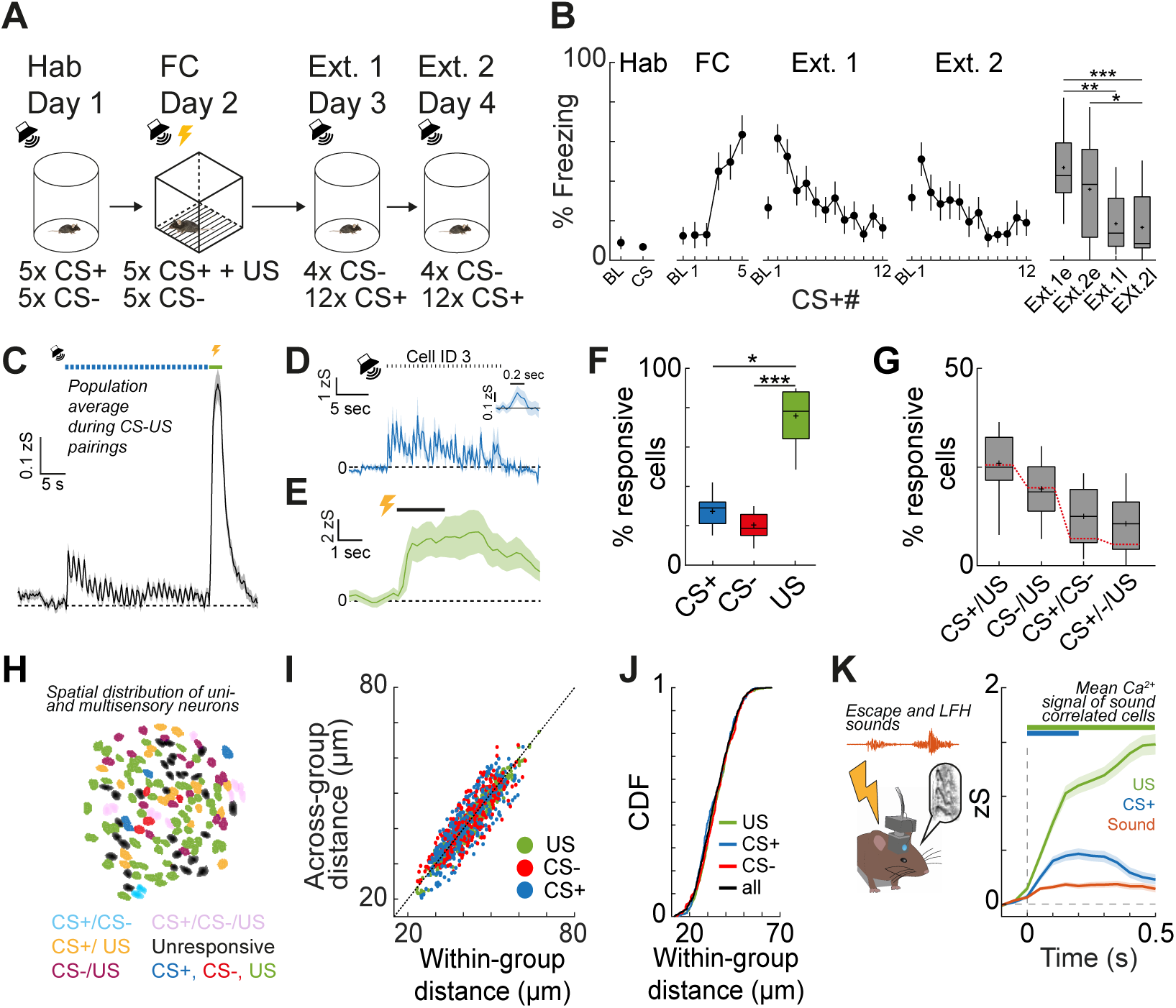
Mixed selectivity tone CS+ and shock US coding of MGB neurons upon fear conditioning. **A)** Details of the four-day fear conditioning paradigm. **B)** CS+ freezing during the habituation, fear conditioning as well as extinction days (Ext.1, Ext.2. e and l indicate early and late phases of extinction, i.e. the first four or last four CS+ of the session. Friedman test, p < 0.001, followed by Dunn’s multiple comparisons test, N = 15 mice). **C)** Mean population response of one example animal to the CS+ and US. Blue dots indicate CS+ tone pips. Green bar indicates shock US. **D, E)** Example cell response to the CS+ (D) and US (E). Mean ± s.e.m. of five trials. Dots indicate CS+ tone pips. Inset represents average response to single pips. **F)** Proportion of CS+, CS- and US responsive neurons. Friedman test, p < 0.001, followed by Dunn’s multiple comparisons test: CS+ vs. US, P < 0.05; CS- vs. US, p < 0.001 (N = 9 mice). **G)** Proportion of mixed selectivity CS+/- and US coding neurons. Red line indicates chance overlap level. **H)** Example spatial map of unisensory and multisensory mixed selectivity CS and US coding neurons in MGB. **I)** Relationship between within response group and across response group pairwise spatial distance between neurons (n = 855 cells, N = 9 mice). **J)** Cumulative distribution function of pairwise distances between all, US-responsive, CS+ and CS-responsive neurons (n = 855 cells, N = 9 mice). **K)** Mean Ca^2+^ activity of sound-correlated neurons during shock evoked sound events (e.g. mouse escape sounds and low frequency harmonic vocalizations (LFH), orange), the first CS+ pip (blue) and the US (green) from n = 550 CS+/US or n = 4956 sound event trials from 110 cells out of N = 3 mice.

To test if US responses are solely driven by movement of the animal or self-induced sounds, e.g. escape runs or low frequency harmonic vocalizations during the aversive foot shock, we correlated the activity of individual MGB neurons with movement speed or the occurrence of sounds in the context. First, a large proportion of MGB neurons exhibited an apparent correlation between movement speed and Ca^2+^ activity during the US. However, this is most likely due to the simultaneous occurrence and conflation of the 2 s aversive foot shock and the behavioural output (escape), given that the activity in the large majority of MGB neurons was not motion or speed correlated during the habituation period (**Figure S3**). Additionally, US responses in MGB cannot be solely explained by the auditory environment, i.e. movement sounds or low frequency harmonic vocalization (Grimsley et al., 2013; Williams et al., 2008) of the animal during the aversive foot shook, given that US and CS+ responses were typically substantially larger in sound correlated neurons than responses to self-evoked sounds of the animal (**Figures 2K and S4**). This indicates that MGB US responses are most likely driven by direct somatosensory input, pain signals or aversive state switches.

In summary, our data demonstrates that auditory thalamus neurons are strongly responsive to both pure auditory tones as well as aversive stimuli. This integration of CS and US inputs underlines that MGB neurons are ideal candidates for sensory plasticity upon associative learning (Bordi and LeDoux, 1994a; Weinberger, 2011).

### Neural response dynamics of MGB neurons upon fear conditioning

MGB neurons, particularly in the medial subdivision, have been shown to potentiate auditory CS responses upon fear learning (Bordi and LeDoux, 1994b; Ledoux et al., 1987; Ryugo and Weinberger, 1978). However, their response diversity and dynamics on the population level upon associative learning remain unknown. To understand the learning-related dynamics of MGB neurons, we followed the activity of large populations of individual MGB neurons across the four-day fear conditioning paradigm. Using a cluster analysis approach, we classified CS+ responsive neurons according to their response dynamics before and after fear conditioning and fear extinction (**Figures 3A and 3B**). On the habituation day and on the two extinction days, we identified eight subgroups of CS+ responsive neurons. 12% ± 1% of cells show stable CS+ responses across days. The remainder could be separated in the following subgroups of plastic neurons: neurons that abolish their complete (11% ± 1%, CS down cells) or onset (8% ± 3%, on-down cells) CS+ response after fear conditioning, neurons with enhanced CS+ responses when the animal is in a high fear state (24% ± 4%, fear cells), neurons that are inhibited when the animal is in a high fear state (2% ± 1%, fear-down cells) as well as neurons that enhance or decrease their response when the animal extinguished the fear behaviour (21% ± 5%, extinction-up cells; 14% ± 3%, extinction-down cells). Additionally, we identified cells that had stable, enhanced CS+ responses after fear learning (8% ± 2%, persistent cells, **Figure 3C**). Similar subgroups were found for CS-responsive neurons. However, in contrast to the US-paired CS+, the group of CS-stable cells was most prominent across days (**Figures S5A-S5C**). Additionally, we found similar proportions of CS+ (**Figure S2B**) and CS-responsive (**Figure S2C**) neurons during the habituation, fear conditioning and extinction days. However, the proportion of neurons that were plastic and changed their CS responses across days was significantly higher in the CS+ group (88% ± 1%) compared to the CS-group (60% ± 7%, 2-way ANOVA, p < 0.0001, followed by Tukey post hoc test p < 0.05), while the proportion of stable neurons was higher in the CS- (40% ± 7%) compared to the CS+ group (12% ± 1%, 2-way ANOVA, p < 0.0001, followed by Tukey post hoc test, p < 0.05, **Figure 3D**), indicating that neural response plasticity is learning specific and more prominent for the paired conditioned stimulus than the control stimulus.

**Figure 3:**
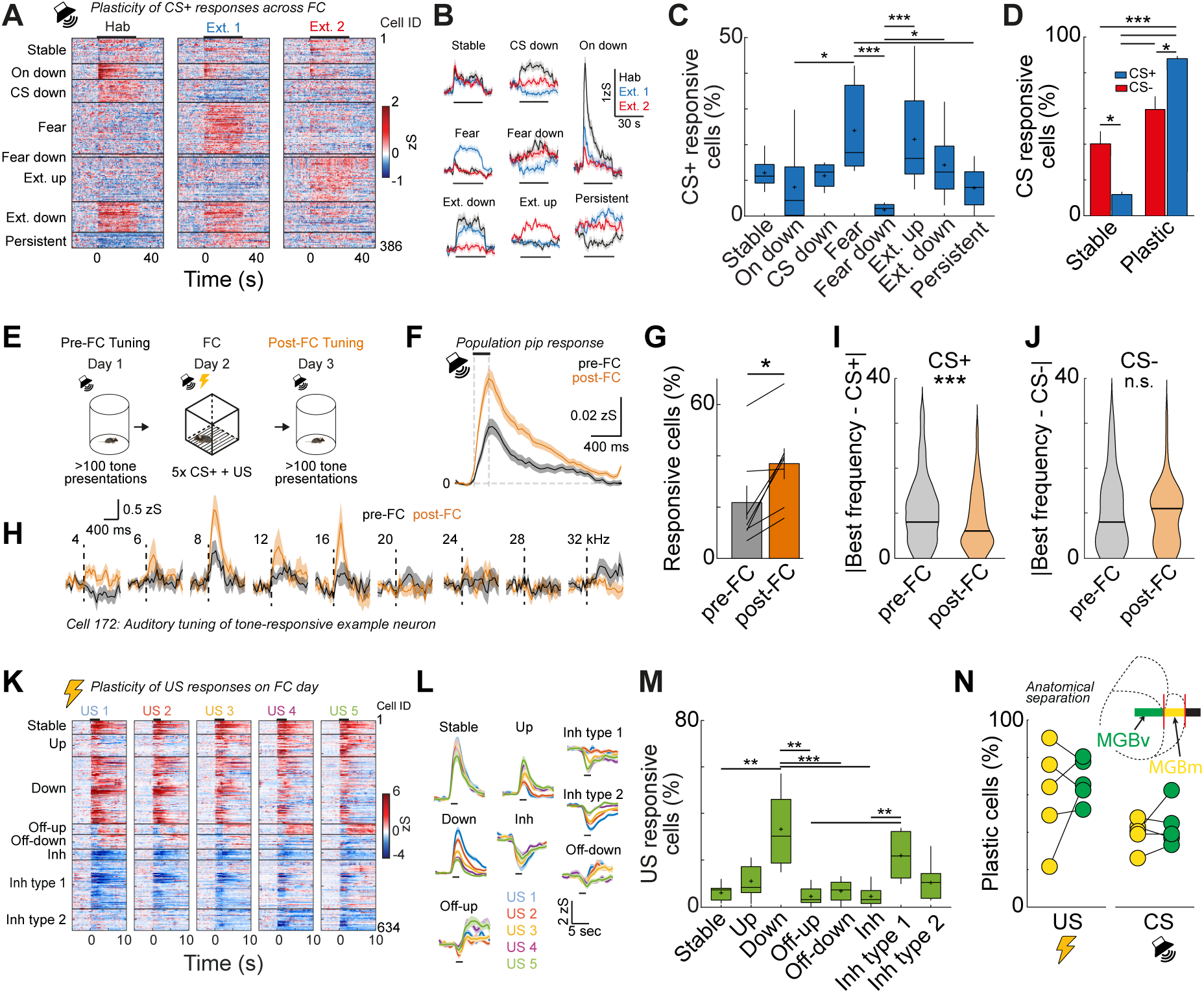
Single cell response plasticity in MGB upon fear learning. **A)** Heat map of single cell CS+ responses on the habituation, extinction 1 and extinction 2 days. Cells were clustered into groups depending on their CS+ response pattern (n = 386 cells, N = 9 mice). **B)** Average traces of neuronal clusters in A. **C)** Proportion of individual plasticity groups within CS+ responsive cells / animal (Kruskal-Wallis test, p < 0.05, followed by Dunn’s multiple comparisons test; On down vs. fear, p < 0.05; fear vs fear down, p < 0.001; fear vs persistent, p < 0.05; fear down vs extinction up, p < 0.001; fear down vs extinction down, p < 0.05; N = 9 mice). **D)** Proportion of CS+ and CS-stable and plastic neurons (2-way ANOVA followed by Tukey post hoc test, p < 0.001; stable CS- vs. stable CS+, p < 0.05; stable CS- vs. plastic CS-, p < 0.05; stable CS- vs. plastic CS+, p < 0.001; stable CS+ vs. plastic CS-, p < 0.001; stable CS+ vs. plastic CS+, p < 0.001; plastic CS- vs. plastic CS+, p < 0.05). **E)** Experimental paradigm. Auditory tuning was tested before and after fear conditioning. **F)** Population response to individual 200 ms pips before (black) and after (orange) fear conditioning (n = 681 cells, N = 7 mice). **G)** Proportion of tone-responsive cells before and after fear conditioning (Wilcoxon signed rank test, p < 0.05, N = 7 mice). Lines represent individual animals. **H)** Auditory responses of one example neuron before (black) and after (orange) fear conditioning to different tone frequencies (numbers indicate kHz). **I)** Population statistics for BF tuning towards the CS+ (Wilcoxon signed rank test, p < 0.001, n = 284 neurons, 7 mice). Horizontal lines represent median. **J)** Population statistics for BF tuning towards the CS- (Wilcoxon signed rank test, p > 0.5, n = 284 neurons, 7 mice). Horizontal lines represent median. **K)** Heat maps of single cell US responses to the five US stimulations during the fear conditioning day (n = 634 cells, N = 9 mice). **L)** Average response of plasticity subtypes of US-responsive MGB neurons (n = 634 cells, N = 9 mice, see also Figure S3). **M)** Proportion of individual plasticity groups within US responsive cells / animal (Kruskal-Wallis test, p < 0.01 followed by Dunn’s multiple comparisons test; Stable vs down, p < 0.01; down vs off-up, p < 0.001; down vs. off-down, p < 0.01; down vs. inh, p < 0.001; off-up vs inh type 1, p < 0.01; inh vs inh type 1, p < 0.01; N = 9 mice). **N)** Proportion of US (N = 5 mice, p > 0.05, Wilcoxon signed-rank test) and CS plastic cells (N = 5 mice, p > 0.05, Wilcoxon signed-rank test) in the ventral (MGBv) and medial (MGBm) subdivisions of MGB. Insert: Schematic of GRIN lens location above the different MGB subdivisions.

All-in-all, this data reveals a broad responses diversity of MGB neurons upon fear learning that extends previous observations of fear conditioning potentiated neurons (Edeline and Weinberger, 1992; Maren et al., 2001; Ryugo and Weinberger, 1978). The plastic CS+ subgroups are similar to previously described functional groups in the amygdala (Amano et al., 2011; Grewe et al., 2017; Gründemann et al., 2019; Herry et al., 2008). While fear and extinction neurons are the most prominent groups, they appear alongside other distinct subgroups, indicating that diverse CS+ response plasticity occurs not only in the amygdala, but also upstream in auditory thalamus.

MGB neurons are diversely tuned to auditory frequencies (Bordi and LeDoux, 1994a) and individual neurons were reported to change their frequency tuning upon associative learning (Edeline and Weinberger, 1991b, 1991a, 1992). To estimate changes in auditory frequency tuning in large populations of individual MGB neurons before and after fear conditioning, we presented 200 ms pips of at least eleven different pure tone frequencies (1 - 40 kHz) at 65 - 85 dB to freely moving mice while simultaneously imaging MGB Ca^2+^ activity (**Figure 3E**). We found that the mean pip response of the MGB population across all frequencies was nearly doubled after fear conditioning (**Figure 3F**). Furthermore, the proportion of tone responsive neurons was increased post conditioning (**Figure 3G**, before: 22% ± 7%, after: 37% ± 6%, Wilcoxon signed-rank test, p < 0.05, N = 7 mice). Besides a general enhancement of pip responses across the whole population and all tones, we found that fear conditioning induces a specific enhancement of selective frequencies compared to the pre-conditioning state (**Figure 3H**), which resulted in a shift of the best frequency towards the conditioned stimulus (**Figure 3I**, pre FC: |Best frequency - CS+| = 11 ± 0.45 kHz, post FC: |Best frequency - CS+| = 8 ± 0.39 kHz, Wilcoxon signed-rank test, p < 0.001, n = 284 from N = 7 mice). This shift was specific for the CS+ and did not occur for the CS- (**Figure 3J**, p > 0.05, Wilcoxon signed-rank test). This data indicates that auditory fear conditioning affects the auditory frequency tuning of MGB neurons in a stimulus specific manner. However, the absolute shift across the population is small (ca. 3 kHz), suggesting that MGB preserves a broad tuning range for reliable sensory representation, despite a stronger representation of the CS+.

In addition to plasticity of CS tone representations, we found that US responsive neurons can be subdivided into stable (6% ± 1%) and plastic cells (94% ± 1%) during fear conditioning. The majority of US responsive cells was plastic and exhibited dynamic intra-session representations of the US (Mann-Whitney test, p < 0.001). Using a similar cluster analysis approach, we identified neurons that demonstrated intra-session potentiation or depression and subtypes of inhibited as well as off-responsive neurons that potentially signal relief from the shock (**Figures 3K-3L**). However, despite its prominent diversity the US response type was not predictive of CS plasticity in MGB neurons. US response and plasticity type did not overlap with CS response and plasticity type above chance levels (**Figure S6**), indicating that US inputs *per se* do not drive plasticity in MGB neurons. Nevertheless, adaptive US responses in MGB could act as an upstream teaching signal in addition to local circuit mechanisms (Krabbe et al., 2019), which direct plasticity in downstream areas like the amygdala.

MGB is subdivided into a first order, auditory cortex-projecting nucleus (MGBv) as well as higher order nuclei (MGBd, MGBm), which send axons to cortical and limbic brain areas, e.g. the amygdala (Doron and LeDoux, 1999). To test if MGB CS and US plasticity types are different between first order and higher order nuclei, we subdivided the cells depending on their location in the GRIN lens field of view between MGBv and MGBm after anatomical verification of the lens front location for all mice where MGBv and MGBm were simultaneously imaged (N = 5 mice, **Figures S7A and S7B**). Similar to the total population of MGB neurons, large fractions of MGBv and MGBm neurons exhibited plasticity to the US or CS+ (**Figure 3N**). Nevertheless, plastic cells were not significantly different between either subdivision (**Figure 3N**) and the diversity of plasticity subtypes was similar in the first order (MGBv) vs. higher order (MGBm) area of auditory thalamus (**Figures S7C-S7E**).

Overall, we found that responses of individual MGB neurons to the CS and US are plastic upon fear conditioning. In addition to previously reported potentiated auditory neurons, we find highly diverse subtypes of CS or US plastic neurons that go beyond FC-driven response potentiation and are distributed similarly across both first order and higher order MGB subdivisions. US response subtypes were not predictive of CS plasticity, nor were the proportions of subgroups or their enrichment predictive of behavioural outcomes on an animal-by-animal basis (**Table 1**), indicating that MGB neurons might play diverse roles in guiding memory formation during associative learning.

**Table 1:** Correlation between neuronal activity and freezing. Significant correlations are labelled in red.

| CS+ clusters/CS+ Freezing | Stable | On down | CS down | Fear | Fear Down | Extinction up | Extinction down | Persistent |
| --- | --- | --- | --- | --- | --- | --- | --- | --- |
| X1 early | $R^2 = 0.0667 P > 0.05$ | $R^2 = 0.0004 P > 0.05$ | $R^2 = 0.396 P > 0.05$ | $R^2 = 0.0044 P > 0.05$ | $R^2 = 0.0040 P > 0.05$ | $R^2 = 0.0001 P > 0.05$ | $R^2 = 0.0097 P > 0.05$ | $R^2 = 0.2338 P > 0.05$ |
| X2 late | $R^2 = 0.0013 P > 0.05$ | $R^2 = 0.0546 P > 0.05$ | $R^2 = 0.0011 P > 0.05$ | $R^2 = 0.0521 P > 0.05$ | $R^2 = 0.0866 P > 0.05$ | $R^2 = 0.0684 P > 0.05$ | $R^2 = 0.0036 P > 0.05$ | $R^2 = 0.0361 P > 0.05$ |

| US clusters/CS+ Freezing | Stable | Up | Down | Off-up | Off down | Inh | Inh type 1 | Inh type 2 |
| --- | --- | --- | --- | --- | --- | --- | --- | --- |
| X1 early | $R^2 = 0.0078 P > 0.05$ | $R^2 = 0.2738 P < 0.05$ | $R^2 = 0.0041 P > 0.05$ | $R^2 = 0.2566 P > 0.05$ | $R^2 = 0.2178 P > 0.05$ | $R^2 = 0.0064 P > 0.05$ | $R^2 = 0.0094 P > 0.05$ | $R^2 = 0.0147 P > 0.05$ |
| X2 late | $R^2 = 0.0031 P > 0.05$ | $R^2 = 0.0165 P > 0.05$ | $R^2 = 0.0306 P > 0.05$ | $R^2 = 0.0258 P > 0.05$ | $R^2 = 0.0248 P > 0.05$ | $R^2 = 0.00305 P > 0.05$ | $R^2 = 0.0087 P > 0.05$ | $R^2 = 0.0119 P > 0.05$ |

| CS- clusters/CS+ Freezing | Stable | CS down | Fear | Fear Down | Extinction up | Extinction down | Persistent |
| --- | --- | --- | --- | --- | --- | --- | --- |
| X1 early | $R^2 = 0.0920 P > 0.05$ | $R^2 = 0.2272 P > 0.05$ | $R^2 = 0.0010 P > 0.05$ | $R^2 = 0.0627 P > 0.05$ | $R^2 = 0.0634 P > 0.05$ | $R^2 = 0.00390 P > 0.05$ | $R^2 = 0.0770 P > 0.05$ |
| X2 late | $R^2 = 0.0309 P > 0.05$ | $R^2 = 0.0147 P > 0.05$ | $R^2 = 0.0238 P > 0.05$ | $R^2 = 0.1769 P > 0.05$ | $R^2 = 0.0581 P > 0.05$ | $R^2 = 0.0280 P > 0.05$ | $R^2 = 0.3992 P > 0.05$ |

Correlation neuronal activity and CS- /CS+ freezing
| CS+ clusters | Stable | On down | CS down | Fear | Fear Down | Extinction up | Extinction down | Persistent |
| --- | --- | --- | --- | --- | --- | --- | --- | --- |
| CS-/CS+ X1 early | $R^2 = 0.0080 P > 0.05$ | $R^2 = 0.1112 P > 0.05$ | $R^2 = 0.0156 P > 0.05$ | $R^2 = 0.0025 P > 0.05$ | $R^2 = 0.0134 P > 0.05$ | $R^2 = 0.0027 P > 0.05$ | $R^2 = 0.0938 P > 0.05$ | $R^2 = 0.0399 P > 0.05$ |

| US clusters | Stable | Up | Down | Off-up | Off down | Inh | Inh type 1 | Inh type 2 |
| --- | --- | --- | --- | --- | --- | --- | --- | --- |
| CS-/CS+ X1 early | $R^2 = 0.0113 P > 0.05$ | $R^2 = 0.1444 P < 0.05$ | $R^2 = 0.0059 P > 0.05$ | $R^2 = 0.4772 P < 0.01$ | $R^2 = 0.1439 P > 0.05$ | $R^2 = 0.0408 P > 0.05$ | $R^2 = 0.0128 P > 0.05$ | $R^2 = 0.0347 P > 0.05$ |

**Table 1: Correlation between neuronal activity and freezing.**
| CS- Clusters | Stable | CS down | Fear | Fear Down | Extinction up | Extinction down | Persistent |
| --- | --- | --- | --- | --- | --- | --- | --- |
| CS-/CS+ X1 early | $R^2 = 0.0004 P > 0.05$ | $R^2 = 0.4702 P < 0.01$ | $R^2 = 0.0010 P > 0.05$ | $R^2 = 0.0056 P > 0.05$ | $R^2 = 0.1256 P > 0.05$ | $R^2 = 0.0004 P > 0.05$ | $R^2 = 0.1068 P > 0.05$ |

### Amygdala-projecting MGB neurons are plastic upon associative fear learning

Higher order MGBm neurons project to different output targets including primary and secondary auditory cortex, striatum and the basolateral amygdala (Chen et al., 2019; Kimura et al., 2003; Kwon et al., 2014; Smith et al., 2019). Enrichment of plastic neurons in the MGB→BLA pathway might be crucial for fear learning, given BLA’s key role in aversive memory formation (LeDoux, 2000). Similar to previous reports (Doron and LeDoux, 1999), we found BLA-projecting MGB neurons to be typically located in higher order MGB areas and particularly enriched in the medial subdivision of MGB (**Figures 4A and 4B**, MGBm, 70% ± 7% vs. 23% ± 6% dorsal, MGBd or 7% ± 3% ventral, MGBv). By-and-large, 30% of MGBm and 10% of MGBd neurons were amygdala-projecting (**Figure 4C**). In contrast, only a small fraction of MGBv neurons (2%) were retrogradely traced from the amygdala. Furthermore, 85% ± 2% of amygdala-projecting MGB neurons (**Figures 4D and S8A**, N = 6 mice) were positive for the higher-order MGB area marker calretinin (Lu et al., 2009), suggesting that calretinin is a highly prevalent but not exclusive marker of BLA-projecting MGB neurons.

**Figure 4:**
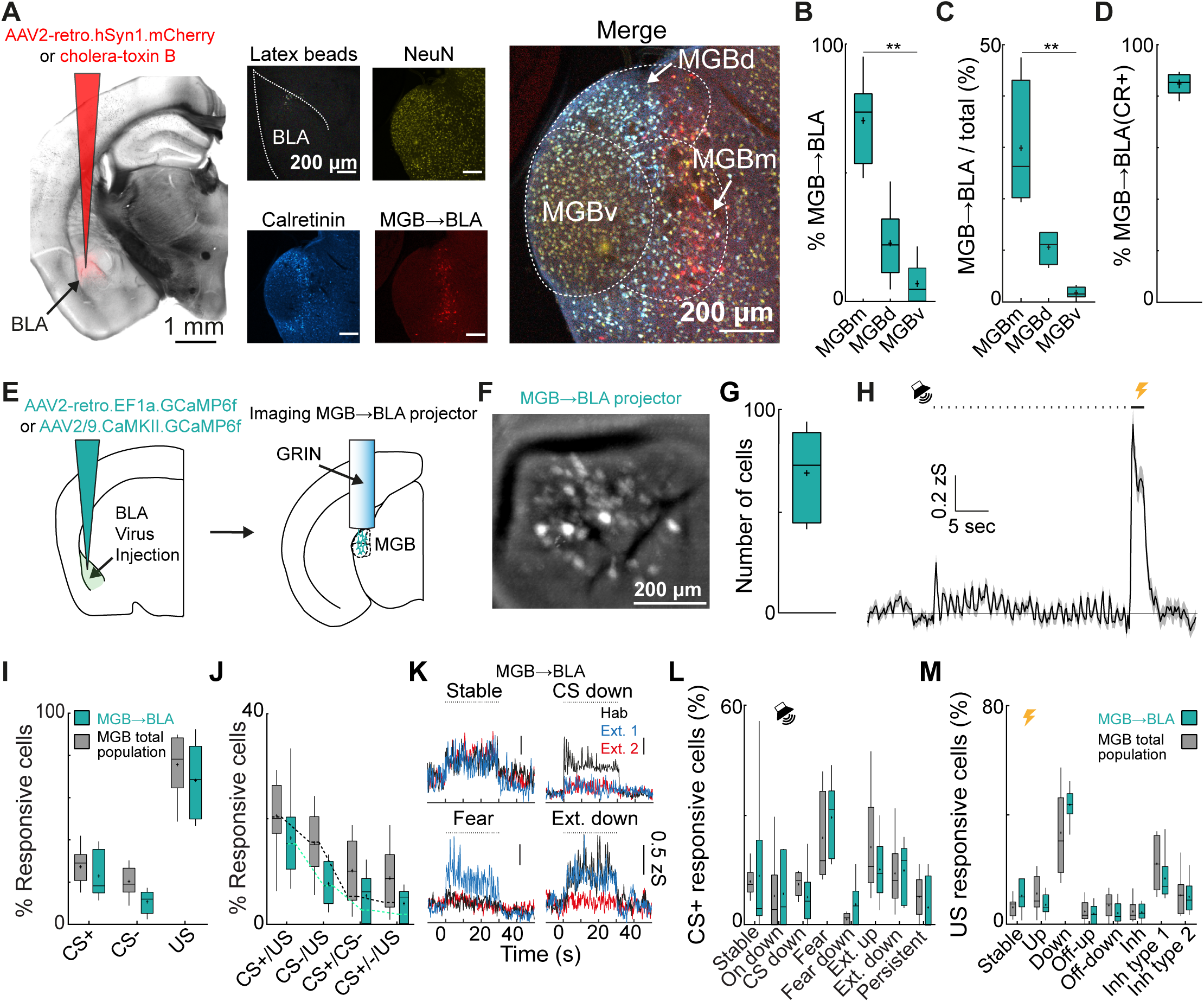
Functional subclasses of CS and US coding neurons are not enriched in amygdala projecting MGB neurons. **A)** Injection of AAV2-retro.hSyn1.mCherry.WPRE.hGHp(A) and latex beads in the basolateral amygdala (BLA). MGB was counterstained for calretinin (cyan) and NeuN (yellow) to quantify the BLA projectors (red). **B)** Distribution of BLA-projecting neurons within MGB (N = 6 mice, Friedman test p < 0.001, followed by Dunn’s multiple comparisons test %MGBm vs. %MGBv, p < 0.01). **C)** Region-specific proportion of BLA-projecting neurons within MGB subdivisions (N = 4 mice, Friedman test, p < 0.01, followed by Dunn’s multiple comparisons test, %MGBm vs. %MGBv, p < 0.05). **D)** Proportion of calretinin-positive BLA-projecting neurons. **E)** Schematic of viral strategy and location of GRIN lens in MGB to image neuronal activity of MGB→BLA-projecting neurons. **F)** MGB field of view with MGB→BLA-projecting neurons. **G)** Number of identified individual components per animal (69 ± 9, N = 6 mice). **H)** Mean population response of one example animal to the CS+ and CS-. Black dots indicate CS+ tone pips. Bar indicates shock US. **I)** Proportion of CS+, CS- and US responsive neurons for the total MGB population and amygdala-projecting neurons (2-way ANOVA, main effect group, F_(1,13)_ = 3.3, p > 0.05, N = 9 total MGB population mice and N = 6 MGB →BLA projection neurons mice, see also Figure 2F). **J)** Proportion of mixed selectivity CS+/- and US coding neurons for the total MGB population and amygdala projecting neurons (2-way ANOVA, F_(1,13)_ = 3.9, p > 0.05, N = 9 mice for the total MGB population and N = 6 mice for the population of MGB→BLA projection neurons, see also Figure 2G). Dotted lines indicate chance overlap level. **K)** Examples traces of groups of stable, onset down, fear and extinction neurons. **L)** Proportion of individual plasticity groups within CS+ responsive cells / animal (2-way ANOVA, main effect group, F_(1,13)_ = 1.2, p > 0.05,, N = 9 mice for the total MGB population and N = 6 mice for the population of MGB→BLA projection neurons). **M)** Proportion of individual plasticity groups within US responsive cells / animal (2-way ANOVA, F_(1,13)_ = 0.5, p > 0.05, N = 9 mice for the total MGB population and N = 6 for the population of MGB→BLA projection neurons).

To test if MGB→BLA projection neurons are necessary for fear learning (Antunes and Moita, 2010; Apergis-Schoute et al., 2005; Han et al., 2011; Pereira et al., 2019), we specifically expressed the inhibitory opsin ArchT in MGB→BLA projectors (**Figures S9A-S9D**). Inhibition of MGB→BLA projectors during CS-US pairing on the conditioning day had no effect on fear acquisition (**Figures S9E-S9G**, mean freezing during the last two CS+ presentations, control: 50% ± 5% freezing, ArchT: 45% ± 8% freezing, p > 0.05, Mann-Whitney Test). However, freezing levels were significantly reduced upon fear test 24 h later (**Figure S9H**, mean freezing during the first four CS before extinction, control: 50% ± 6% freezing, ArchT: 27% ± 7% freezing, p < 0.05, Mann-Whitney Test) indicating that activity in MGB→BLA projectors is necessary for the consolidation of fear memories.

To test the physiological function and neuronal activity of amygdala-projecting MGB neurons in fear learning, we used a retrograde virus approach to specifically express GCaMP6f in MGB→BLA-projectors (see Methods, **Figures 4E and S8B**). On average, we could identify 69 ± 9 BLA-projecting GCaMP6f-positive MGB neurons per mouse (**Figures 4F and 4G**, N = 6 mice). Similar to the total MGB population average response (**Figure 2C**), MGB→BLA projectors were activated by the CS+ and the US (**Figure 4H**). Across animals, 22% ± 5%, 10% ± 2% or 68% ± 7% of neurons were responsive to the CS+, CS- or US, respectively. These proportions are comparable to the total MGB population (**Figure 4I**, 2-way ANOVA, p > 0.05). Furthermore, we did not find that combinations of CS+, CS- and US coding neurons were enriched above chance levels (**Figure 4J**). Using a cluster analysis approach, we found the same subgroups of CS+ plasticity types in the subpopulation of MGB→BLA projecting neurons (**Figures 4K and 4L**) across the conditioning paradigm, including stable cells, onset-down cells, CS- down cells, fear cells, fear-inhibited cells, extinction up cells, extinction down cells as well as persistent cells (**Figures S8C-S8E**). The proportions of the CS+ plasticity subgroups were similar to the total population in MGB (2-way ANOVA, p > 0.05). Analogous to the CS+ representation across days, we found comparable proportions of US plasticity types in MGB→BLA projectors when compared to the total population (**Figures 4M and S8F**, 2-way ANOVA, p > 0.05).

This data demonstrates that CS as well as US information is encoded by BLA-projecting MGB neurons, identifying MGB→BLA projectors not only as a source of CS tone inputs but also as a strong source of aversive US signals (see also Bordi and LeDoux, 1994a). However, CS and US plasticity is functionally diverse beyond response potentiation and, compared to the total MGB population, CS and US signalling is not enriched in this specific subpopulation of amygdala-projecting MGB neurons.

### Population coding and representation of the conditioned stimulus across days

Next, we tested if the CS+ and CS- can be decoded from Ca^2+^ activity based on MGB population activity. First, we trained a three-way quadratic decoder to distinguish between CS+, CS- and baseline activity within the same session (see Methods). To balance for different cell population sizes between animals, we randomly sub-selected 40 cells for each animal and averaged decoder accuracy across 50 independent runs. Furthermore, to account for different numbers of CS+ and CS-presentations, we only decoded the first four CS+ and CS-presentations. Within each individual session, the decoders achieved high classification accuracy (**Figure 5A**, > 80% compared to 33% chance level) indicating a distinct representation of the individual CSs by the MGB population. Surprisingly, decoding accuracy was higher in the population of MGB→BLA projectors compared to the total MGB population, except for the fear conditioning day. To test if CS tones can be accurately detected across days from MGB population activity, we next trained sets of two-way decoders to distinguish between baseline and CS+ or CS-responses for each experimental day and tested the trained decoders across days (**Figure 5B**). Strikingly, we found that decoder accuracy is robust across days reaching decoding levels of ca. 70% or higher, for both the CS+ and the CS- (**Figure 5C**). This is in contrast to amygdala population coding, where decoding levels for the CS+ drop to chance levels after fear conditioning (Grewe et al., 2017), indicating that MGB population representations of CS tones are stable despite associative learning. Furthermore, we found a drop in CS+ encoding in MGB→BLA projectors during the fear conditioning day, which recovered afterwards (**Figure 5C**, see also **Figure 5A**), indicating temporary changes in CS+ encoding during associative learning.

**Figure 5:**
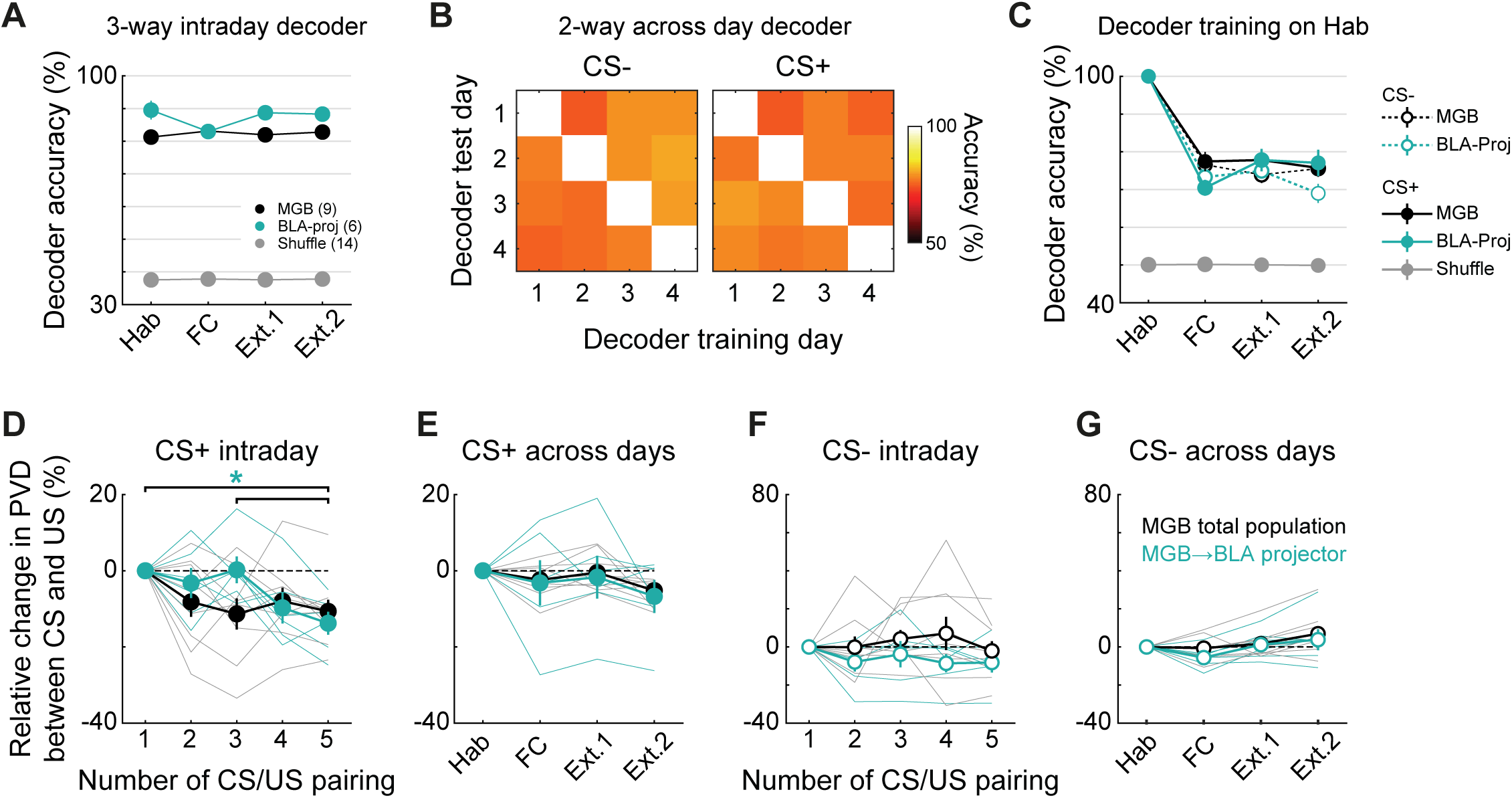
MGB population dynamics are stable across days. **A)** Intraday three-way decoder of CS+, CS- and baseline population responses in CaMKII-positive (black) and identified amygdala-projecting MGB neurons (turquoise) reached a minimum mean accuracy of 81% across animals. Decoder accuracy dropped to chance levels for decoders trained on randomly label training sets. **B)** Intra- and across day accuracy of decoders trained on CS+ or CS- vs. baseline responses, respectively. 1: Hab, 2: FC, 3: Ext. 1, 4: Ext.2. **C)** Quantification of intra and across day decoder accuracy for decoders trained on habituation day data. Mean decoder accuracy across days is > 70 % for CS+ and CS-population responses in CaMKII-positive and identified amygdala-projecting MGB neurons. **D-G)** Relative change in Euclidean population vector distance between the CS+ (D, E) or CS- (F-G) and the US within the fear conditioning session (D,F) or across the individual days of the behavioural paradigm (E,G). Statistics: D: Friedman test across the relative change in CS+ to US PVD of MGB→BLA-projectors (p < 0.01), Dunn-sidak multiple comparisons test 1st and 3rd vs. 5th CS/US pairing, p < 0.05. G: Friedman test across the relative change in CS- to US PVD of the toal BLA-population (p < 0.05), Dunn-sidak multiple comparisons test FC vs. Ext.2: p < 0.05. All other data sets in D-G: p > 0.5. MGB population: N = 9 mice, MGB→BLA-projectors: N = 6 mice.

Finally, we compared the population vector distance (PVD) between the evoked population responses to the CS+ and CS- and the evoked population responses to the US (**Figures 5D-5G**). During fear conditioning, we found a decrease of the PVD between the CS+ and the US with consecutive CS-US pairings (**Figure 5D**) for both the total MGB population as well as for MGB→BLA-projectors. However, the time courses of the PVD change were different and only the MGB→BLA population reached a significant change in PVD at the end of the session indicating different population dynamics for this subgroup of MGB neurons during associative learning (Friedman test, p < 0.01, 1^st^ and 3^rd^ versus 5^th^ paring: Dunn-Sidak multiple comparisons test, p < 0.05). Importantly, no changes were found between the evoked responses to the CS- and US during conditioning. In contrast to previous observations in the amygdala (Grewe et al., 2017), the PVD between the CS+ and the US changes were not preserved post conditioning on the extinction days and the population representation recovered to pre-conditioning levels similar to the observations using two-way or three-way decoders (**Figures 5A and 5C**). However, note that we found a significant difference between the CS- to US PVD of the total MGB population between the conditioning day and the second extinction day (Friedman test, p < 0.05, FC versus Ext. 2: Dunn-Sidak multiple comparisons test, p < 0.05). Nevertheless, we did not find a change in the CS- to US PVD between the habituation, fear conditioning and early extinction days indicating that the CS-representation is stable during high fear states in relation to the habituation day and shows no significant drift intraday during fear conditioning. Furthermore, the CS-drifts further away from the US in comparison to the habituation day which might be reflective of an enhanced safety signal after extinction.

In addition to the general lack of consolidation of population level changes across fear learning, the strength of PVD-changes between the CS+ and US were not predictive of learned freezing behaviour on an animal-by-animal basis (**Figure S10**).

Taken together, this data indicates that high dimensional representations of CS+ tones are stably encoded in MGB populations across associative fear learning, despite plastic changes in single cell response patterns. In contrast to the basolateral amygdala (Grewe et al., 2017), MGB population representations of sensory stimuli only transiently change during associative fear learning and reset overnight, which might be crucial for unbiased representations of stimuli from the environment.

## Discussion

By imaging large populations of MGB neurons, we find that auditory thalamus is a site of diverse neuronal plasticity during associative fear learning on the level of single cells as well as the total MGB population. However, changes in MGB population level coding are only transient and do not consolidate overnight, which might be instructive for plastic changes in downstream structures during learning, yet allows for long-term stability of sensory coding across days.

On the level of individual MGB neurons, we observed associative learning-induced CS+ response potentiation that resembles classic studies which demonstrated that auditory thalamus exhibits enhanced responses to aversive conditioned tones (Edeline et al., 1990; Hennevin et al., 1993; Maren et al., 2001; Ryugo and Weinberger, 1978). However, recording simultaneously from large populations of individual neurons in MGB during fear conditioning, we find diverse plasticity patterns (at least 7) that are similar or go beyond previously reported plasticity types in cortical (Dalmay et al., 2019) or limbic areas (Amano et al., 2011; Grewe et al., 2017; Gründemann et al., 2019; Herry et al., 2008) downstream of MGB. Changes in CS responsiveness of individual MGB neurons upon fear learning and extinction were bidirectional, i.e. potentiated or depressed, and depended on the behavioural state of the animal. For example, we find different functional types that are enhanced or depressed particularly in high fear or extinction states. This extends the notion of unidirectional response potentiation in MGB upon associative learning and demonstrates that auditory thalamus neurons exhibit heterogeneous, adaptive signaling of threat-predicting auditory stimuli of the environment.

Besides auditory stimuli, MGB neurons signal the aversive foot shock (US) during fear conditioning (Bordi and LeDoux, 1994a; Ryugo and Weinberger, 1978). Indeed, we found that the proportion of foot shock encoding neurons exceeds the number of tone CS+ encoding neurons in auditory thalamus of freely moving animals. The large proportion of US encoding neurons in MGB and the strength of the US signal on the population level could not be explained by movement of the animal or self-vocalization induced activation of MGB (Schneider et al., 2018; Šuta et al., 2008). Aversive US responses are considered to be more prominent in higher order auditory thalamus (MGBm, Bordi and LeDoux, 1994a). Our imaging sites covered both first order (MGBv) and higher-order (MGBm) MGB, and US encoding neurons were equally present in both sites. This demonstrates that multisensory encoding is not an exclusive feature of higher-order areas of MGB in freely moving animals, but can also occur in the first order ventral subdivision that projects to auditory cortex, suggesting that auditory thalamus conveys aversive US information to a broad range of cortical and limbic downstream areas during associative fear learning. Strikingly, US responses were heterogeneous across the population of MGB neurons during fear conditioning. Besides stable US responders, we identified several plasticity types of US responsive neurons, including short term facilitating, depressing or off-responsive neurons. This functional diversity of US neurons indicates that first and higher order auditory thalamus can signal distinct types of instructive information, for example adaptive teaching signals for associative fear learning as well as relief or safety signals upon termination of the US. Future studies need to address if and how these non-uniform adaptive MGB US signals are relayed to specific circuits elements in downstream areas like the amygdala or auditory cortex (Abs et al., 2018; Aizenberg et al., 2015; Gillet et al., 2018; Kim et al., 2016; Krabbe et al., 2018; Lee et al., 2017; Letzkus et al., 2011; Lucas et al., 2016; Namburi et al., 2015; Senn et al., 2014).

CS and US coding neurons are spatially intermingled in auditory thalamus and a large fraction of MGB neurons exhibit mixed selectivity for both the CS tone and US foot shock. The convergence of diverse CS and US responses in individual MGB neurons renders auditory thalamus an ideal site for neuronal plasticity in associative learning (Bordi and LeDoux, 1994a; Weinberger, 2011), which is supported by the finding of large numbers of different subgroups of plastic neurons upon fear conditioning. Nevertheless, similar to observations downstream in the basolateral amygdala (Grewe et al., 2017; Gründemann et al., 2019), the convergence of CS and US responses in MGB neurons was not predictive of the response plasticity of a given neuron. Instead, MGB neurons exhibit manifold functional classes and outcomes of CS/US conversion upon learning (e.g. CS/US responsive cells can become potentiated fear cells or CS down cells), suggesting that heterogeneous fear conditioning-induced auditory response plasticity in MGB is most likely governed by multiple cellular or circuit mechanisms of neuronal plasticity. Indeed, we find subsets of neurons in all groups of CS plastic neurons that were not US responsive during FC (see **Figure S6**). This shows that MGB neurons do not necessarily require converging CS and US input (Ryugo and Weinberger, 1978) to drive functional plasticity upon associative learning, arguing for plasticity mechanisms that go beyond classical Hebbian plasticity, which requires coincidence detection on a millisecond timescale (Markram et al., 1997) and might additionally involve slower, neuromodulatory mechanisms (Brzosko et al., 2019; Izhikevich, 2007; Likhtik and Johansen, 2019) similar to amygdala circuits (Grewe et al., 2017; Johansen et al., 2014) or consolidation during sleep (Hennevin et al., 1993).

Furthermore, brain-wide distributed interacting circuit mechanisms could play a role in the formation of single cell plasticity upon associative fear learning, not only in MGB but across multiple fear-related brain areas (Gehrlach et al., 2019; Herry and Johansen, 2014; Herry et al., 2008; Klavir et al., 2013; Likhtik et al., 2014; Maren et al., 2001; Ozawa et al., 2017; Penzo et al., 2015). The detailed computations within this distributed network (Mobbs et al., 2020) and the role of auditory thalamus are poorly understood. Plasticity and adaptive changes in MGB depend on uni- or multi-synapse feedback circuits from distinct brain areas including the amygdala (Aizenberg et al., 2019; Maren et al., 2001) and cortex (He, 2003). Nevertheless, our data supports the notion that learning-induced modifications of neuronal activity in MGB could drive plastic neuronal responses in downstream areas (Weinberger, 2011). This suggests that at least a part of the heterogeneous response plasticity in amygdala or cortex during associative fear learning could be inherited in a feedforward fashion from adaptive changes in the thalamic relay independent of and in addition to local synaptic and circuit mechanisms (Abs et al., 2018; Gillet et al., 2018; Krabbe et al., 2019).

We found similar proportions of CS and US plastic neurons in first order cortex-projecting and higher order areas of MGB. Higher order areas of MGB are more broadly-projecting areas of auditory thalamus and target among others large parts of auditory cortex as well as striatum and amygdala (Chen et al., 2019; Jones, 1998; Kimura et al., 2003; Smith et al., 2019). Given the amygdala’s prominent role in associative fear learning we hypothesized that CS and US plastic neurons are specifically enriched in MGB→BLA projection neurons when compared to the total population including less plastic MGBv neurons (Edeline and Weinberger, 1991b, 1992). Using a retrograde viral approach, we specifically imaged BLA-projecting MGB neurons to distinguish these cells from the general population. Surprisingly, we found that the proportion of plastic neurons was not enhanced in amygdala-projecting neurons and was similar to the proportion of plastic neurons in the total MGB population. This lack of enrichment of neurons with dedicated functions in associative learning suggests that the MGB→BLA pathway is most likely not a labelled line, but that MGB potentially propagates experience-dependent changes of neuronal activity in associative fear learning to a wider brain network, including auditory cortex and striatum (Chen et al., 2019). This is reminiscent of recent findings that heterogenous behaviour-related neural activity of projection neurons of a given brain area is broadcast simultaneously and in parallel to different downstream targets irrespective of the output pathway (Gründemann et al., 2019; Weglage et al., 2020). Future work including simultaneous multi-site recordings (Jun et al., 2017), targeted-activity dependent neural manipulations (Emiliani et al., 2015) and computational neuroscience tools will be required to delineate how this heterogeneous, widely distributed population code is interpreted by different downstream regions (Headley et al., 2019; Mobbs et al., 2020).

Locally, on the level of the auditory thalamus, CS responses could be decoded reliably from the population level responses of the total MGB ensemble. Within a given day of the fear conditioning paradigm, we could train decoders that reliably distinguish between CS+, CS- or baseline activity with high accuracy. Strikingly, we could also train binary decoders that accurately classified baseline vs. CS+ or CS-presentations across all experimental sessions and along associative fear learning. This suggests that MGB ensembles exhibit stable population level tone representation across days, despite the plasticity of CS responses of individual cells during fear conditioning, which can be stable over weeks (Edeline et al., 1990). This data is supported by the observation that the MGB population vector difference between the CS+ and US decreases during fear conditioning, yet recovers to baseline levels on the next day after the conditioning session. Thus, MGB exhibits stable tone representations on the population level across associative learning, which will be crucial for reliable representations of sensory stimuli from the environment, for example, in light of changing stimulus statistics (Blackwell et al., 2016) in complex environments and plastic single cell responses (see above) or behaviour-driven changes in response amplitudes (Williamson et al., 2015). This is in stark contrast to fear-biased population level changes in sensory representation in the amygdala that are further stabilized and consolidated after learning, and prevent the decoding of tone responses across fear conditioning (Grewe et al., 2017). While the population code of the amygdala stabilizes “fear hijacking” of the sensory representation, MGB exhibits transient changes in population level encoding, which provides a clean slate for future perception that is unaffected by a valence bias. Nevertheless, the transient population level changes during fear conditioning in MGB might be crucial to guide long term population level changes in the amygdala or other downstream areas upon associative learning (Chen et al., 2019; Grewe et al., 2017; Zhang and Li, 2018).

Taken together, our data indicates that auditory thalamus is ideally positioned to exhibit a complex role in guiding neuronal plasticity and valence assignment during associative learning that goes beyond the classical role of auditory processing and response potentiation during conditioning and potentially extrapolates to a broad set of behavioural functions (Gilad et al., 2020). Delineating the neural circuit mechanisms that underlie these highly dynamic representations of uni- and multisensory stimuli in MGB and their experience-dependent plasticity will open new avenues to understand the role of early, pre-cortical sensory relays like auditory thalamus in the formation of sensory percepts and memories that mediate complex behaviours.

## Acknowledgements

We thank Georg Keller and Daniela Gerosa Erni for virus production and V. Jayaraman, R. Kerr, D. Kim, L. Looger, K. Svoboda and the HHMI Janelia GENIE Project for making GCaMP6 available, Ed Boyden and MIT for making ArchT available as well as Alla Karpova and David Schaffer who gifted the rAAV2-retro helper (Addgene plasmid # 81070V). We thank Raymond Strittmatter and Robert Häring for mechanical workshop and electrical engineering support. We thank Yael Bitterman, Sabine Krabbe, Andreas Lüthi and the Gründemann lab for comments on the manuscript. Funding: Research was supported by the Swiss National Science Foundation (Ambizione Fellowship to J.G., SNF Professorship to J.G.), European Research Council (Starting Grant, J.G.), the Forschungsfonds Nachwuchsforschende (M.H.) and the Department of Biomedicine at the University of Basel.

## Author contributions

Conceptualization, J.A.T., M.H. and J.G.; Methodology and imaging experiments, J.A.T., M.H. and J.G.; Experimental support, J.A.F., D.A.G., M.T. and T.L.; Analysis, J.A.T., M.H., C.B. and J.G.; Original draft, J.A.T. and J.G.; Revisions, all authors; Supervision and funding, J.G.

## Declaration of interests

The authors declare no conflict of interest.

## Methods

### Animals

8 to 11-week-old male C57Bl/6J mice were used throughout the study. All experiments were done in accordance with institutional guidelines and were approved by the Cantonal Veterinary Office of Basel-Stadt, Switzerland. Animals were housed on a 12-hour light / dark cycle and food and water were provided ad libitum.

### Surgeries, virus injection and GRIN lens as well as optical fibre implantation

Virus was injected with the help of a stereotaxic apparatus (Kopf Instruments) in the medial geniculate body (for imaging experiments 500 nl, AAV2/5.CaMKII.GCaMP6f.WPRE.SV40, Penn Vector Core, for optogenetic experiments 500 nl AAV2/5.CAG.flex.ArchT.GFP, UNC Vector Core, or AAV2/5.CaMKII.EGFP.WPRE.hGHp(A), VVF Zürich; coordinates: AP: −3.28, ML: −1.9, DV: −3.1 mm) or basolateral amygdala (300 nl, AAV2/9.CaMKII.GCaMP6f.WPRE.SV40, Penn Vector Core; rAAV2-retro.EF1a.GCaMP6f.WPRE, Georg Keller, FMI Vector Core, Basel, Switzerland, for tracing experiments 300 nl, rAAV2-retro.hSyn1.mCherry.WPRE.hGHp(A), VVF Zurich; or 50 nl CTB 555, Invitrogen, for optogenetic experiments 300 nl rAAV2-retro.hSyn1.mCherry.icre.WPRE.hGHp(A), VVF Zürich; coordinates AP: −1.7, ML: −3.6, DV: −3.6 mm) with a glass pipette and a pressure ejection system (Picospritzer) under isoflurane anaesthesia (1 - 2%) and buprenorphine (0.1 mg/kg) and ropivacaine (65 mg/kg) analgesia. The rAAV2-retro helper was a gift from Alla Karpova & David Schaffer (Addgene plasmid #81070). Virus for retrograde labelling of MGB neurons was supplemented with blue non-retrograde polymer microspheres (1:2400, Duke Scientific Corp.) to label BLA injection sites. For miniature microscope experiments, one week after virus injection, a gradient refractive index (GRIN) lens (0.5 or 0.6 mm diameter, Inscopix) was implanted during a second surgery (anaesthesia and analgesia see above). A 0.8 mm diameter craniotomy was drilled above the MGB and a small track was cut with a 0.7 mm sterile needle. The GRIN lens was then slowly advanced into the brain (coordinates: AP: −3.28, ML: −1.9, DV: −3.0 mm), fixed to the skull with light curable glue (Loctite 4305, Henkel) and the skull was sealed with Scotchbond (3M), Vetbond (3M) and dental acrylic (Paladur, Kulzer). A titanium head bar (custom made) was attached to fix the animal during the miniature microscope base plate mounting procedure. For optogenetic experiments, virus (see above) was injected bliaterally in the basolateral amygdala and medial geniculate body as described above. One week later, optical fibres (0.4 mm, 0.5 NA, Thorlabs) were implanted bilaterally above the medial geniculate body (Coordinates: AP: −3.28, ML: −1.9, DV: −2.9 mm). Optical fibres, the wound and skull were fixed and sealed in a similar manner to GRIN lens implantations. Animals were provided with analgesia (buprenorphine, ropivacaine) and their well-being was monitored throughout the entire experimental period.

### Behavioural paradigms and analysis

Behavioural experiments were performed during the animal’s light period. A four-day auditory fear conditioning paradigm was performed in a habituation / test context (days 1, 3, 4) and a fear conditioning context (day 2). Mice were presented with 5 intermingled CS+ and CS-during habituation (6 kHz and 12 kHz, intermingled) in a round plexiglass context. CSs were composed of 27 tone pips (200 ms, 75 dB) presented at a rate of 1.1 Hz (Tucker Davis Technologies, TDT 78 or RZ6). Fear conditioning was performed in a ca. 25 cm square plexiglass box and a shock grid floor (Coulbourn, Noldus). The CS+ (6 kHz and 12 kHz, counterbalanced) was terminated by a 2 s 0.65 mA foot shock 1.1 s after the last tone pip. During the extinction sessions (day 3 and day 4, habituation context), 4 CS- and 12 CS+ were presented. For optogenetic experiments, animals were habituated to the optical fibre attachment procedure for three days before the start of the fear conditioning paradigm. On the fear conditioning day, optical fibres were attached to the optical fibre implant via a ceramic mating sleeve (Thorlabs). MGB ArchT-expressing neurons were continuously inhibited during the five CS- US pairings (starting 2 s before CS onset until 2 s after US offset) with a 565 nm LEDs (M565D2, Thorlabs). Optical stimulation was controlled with a custom-built stimulation setup consisting of an Arduino board (Arduino Uno REV3, Arduino) and LED drivers (LEDD1B, Thorlabs). The light intensity measured at the optical fibre tip was 19 mW and optical fibre implants had a typical attenuation of 30%. Optogenetic experiments were performed and analyzed in a blinded fashion. Behavioural experiments were performed and analysed using Cineplex 3.4.1 (Plexon Inc) or Ethovision 14 (Noldus). Behavioural tracking based on the center of mass of the mouse was performed using inbuilt functions of Cineplex and Ethovision. Freezing was initially detected automatically based on periods of absence of movement (threshold: 1 s) and then manually controlled and adjusted for non-freezing episodes (e.g. grooming) *post hoc*. Plasticity of auditory tuning of MGB neurons was tested with a three-day paradigm. On day one, the animals were exposed to 165 - 195 presentations of 200 ms pure tone pips ranging from 1 - 40 kHz at 65, 75 and 85 dB. Pure tones were presented as a series of three pips at a frequency of 0.5 Hz in a round plexiglass context. The different combinations of tone frequency and sound pressure levels were presented as randomized trials (five repetitions per combination) with a minimum intertrial-interval of 11 s. On the consecutive day, the animals underwent a fear conditioning paradigm as described above (CS frequencies: 8, 16, 20 kHz). On the post-learning test day (day three), mice were exposed to the same presentation of pure tone pips as on day 1. MGB neural activity was imaged throughout the four-day fear conditioning and three-day auditory tuning paradigm using a miniature microscope.

### Miniature microscope imaging

The miniature microscope (nVista2.0, Inscopix) was fixed to the base plate on the mouse’s head before the experiment using head-fixation at the head bar on a flying saucer style running wheel. Mice were initially habituated to this procedure. MGB Ca^2+^ fluorescence was imaged continuously during the behavioural session with the following settings (nVista Software Version: 2.0.4): Framerate: 20 Hz, LED-Power: 50% - 70%, Gain: 1.0-2.0, Image size: 1024 x 1024 or 1080 x 1080 pxl. LED power and gain were adjusted according to GCaMP expression levels and the same settings were used across days for individual mice.

### Image analysis

Raw image data was analysed as previously described (Grewe et al., 2017; Gründemann et al., 2019; Mukamel et al., 2009). Briefly, movies from all behavioural sessions were spatially down sampled (2x), bandpass filtered (Fourier transform) and normalized by the filtered image (ImageJ). The movies were then concatenated and motion corrected using Turboreg (three rounds, Thévenaz et al., 1998). Only movies that motion-corrected across days with final spatial dislocations of < 2 μm were used for Ca^2+^ trace extraction. Principal and independent component analysis-based detection of individual regions of interest (ROIs) was performed on downsampled (5 Hz) ΔF/F movies. ROIs were truncated at 50% peak intensity and limited to a size of 30 pixels (ca. 60 μm). ROIs were initially oversampled (300 ICs) and then overlaid with the maximum intensity projection of the four-day movie. ROIs that did not match individual neurons were discarded. We typically retained 97 ± 5 ICAs per animal for CaMKII-GCaMP6f, N = 19 mice, and 69 ± 9 ICAs per animal for rAAV2-retro.EF1a.GCaMP6f, N = 6 mice. These ICs were then applied to the 20 Hz motion corrected raw fluorescence movie to extract single cell Ca^2+^ traces for further processing.

### Ca^2+^ data analysis

All analysis was based on linearly detrended and z-scored Ca^2+^ traces of individual neurons. Ca^2+^ traces were baselined to the time periods preceding CS or US onset. To identify CS- and US-responsive neurons and their plasticity types across days, 30 s CS and 2.8 s US responses were analysed using a combined statistical and supervised cluster analysis approach as previously described (Gründemann et al., 2019).

Auditory tuning curves were calculated based on the mean Ca^2+^ response during the 250 ms time window after pip onset. Cells were classified as tone-responsive to individual frequency pips if their mean response exceeded 0.5 zS for at least two of the frequencies tested. The best frequency (BF) of a neuron was defined as the frequency that prompted the maximal Ca^2+^ response averaged across trials. The difference in BF to the CS+ for comparison across fear conditioning is calculated on an animal-by-animal basis in absolute values as ΔBF = |BF - CS+|.

CS+, CS- and baseline responses were decoded from MGB Ca^2+^ activity by fitting three-way (CS+ vs. CS- vs. baseline) or two-way (CS+ vs. baseline or CS- vs. baseline) quadratic discriminant analysis classifiers. We classified CS+ and CS-responses based on the first four presentations to balance for uneven numbers of CS+/- presentations across habituation, fear conditioning and extinction days. Baseline responses were sampled from the 30 s periods preceding the CS+ and CS-. Classifiers were trained on the mean response of five consecutive pip responses within one CS (or the baseline period), such that each training set contained 20 input variables per condition (i.e. 40 for two-way decoders and 60 for three-way decoders). The mean response was calculated based on a 300 ms time window after pip onset and classifiers were trained using a 10-fold cross-validation procedure. Decoder accuracy was calculated as the mean of the diagonal of the confusion matrix. Classifiers were trained for each individual animal and are presented as mean decoding accuracy across animals. To balance for unequal cell numbers between the different animals, we randomly selected 40 neurons from each animal and calculated the mean accuracy from 50 independent runs. The population vector distance (PVD) between CS and US responses was calculated based on binned (0.275 s bins to accommodate the 1.1 Hz pip frequency) 30 s CS and 4 s US responses. PVD was calculated as the Euclidean distance between each CS bin and the mean binned US response and then averaged for each 30 s CS. Intraday PVD changes were normalized to the PVD of the first CS and across day PVD changes were calculated as the mean intraday PVD change for all CSs and normalized to the mean PVD of the habituation day.

### Sound recordings and analysis

Acoustic signals of the fear conditioning context were recorded with a PCB Precision Condenser Microphone (Model 377C01) microphone and a RZ6 Auditory Processor (Tucker-Davis Technologies) 50 cm above the fear conditioning context at 195 kHz simultaneous to miniature microscope imaging during the fear conditioning session. Sound waves were high-pass filtered at 1 kHz and spectrograms were computed using short-time Fourier transforms (*spectrogram* function, Signal Processing Toolbox, MATLAB, Mathworks). To detect sound-level correlated neuronal activity, the cross-correlation coefficient between the squared acoustic signal binned in 50 ms and the corresponding Ca^2+^ signal of individual neurons was computed with a maximum lag of 500 ms. Cells were classified as sound-correlated if they exceeded a maximal cross-correlation coefficient of 0.2. Acoustic event onsets were detected based on the peak of the differentiated squared and binned sound wave. Acoustic events were not distinguished between animal movement-related sounds and vocalizations of the animal.

### Histology

After completion of the behavioural experiment, mice were transcardially perfused with ca. 5 ml phosphate buffered saline (PBS, ThermoFisher) followed by 40 ml 4% paraformaldehyde (PFA) in PBS (pH = 7.4). Brains were removed and stored overnight in 4% PFA. 150 μm coronal slices were prepared using a vibratome (Campden Instruments) and immunostained for calretinin using the following solutions and protocol: carrier solution: 1% normal horse serum (NHS, Vector Laboratories) with 0.5% Triton (ThermoFisher) in phosphate buffered saline (PBS, ThermoFisher), blocking solution: 10% NHS with 0.5% Triton in PBS. After several rounds of PBS washes, slices were blocked for two hours at room temperature and incubated in primary antibody in carrier solution (goat anti-calretinin, 1:1000, Swant; rabbit anti-NeuN, 1:3000, Abcam; rabbit anti-GABA, 1:500, SigmaAldrich) overnight at 4 °C. Slices were washed again in PBS and incubated for 2 h at room temperature in secondary antibody in carrier solution (donkey anti-goat 647, 1:1000, ThermoFisher; donkey anti-rabbit 405, 1:1000, Abcam; donkey anti-rabbit 555, 1:1000, ThermoFisher). After four final washes, slices were mounted on slides and cover slipped using 22 x 50 mm, 0.16 - 0.19 mm thick cover glass (FisherScientific). Images were acquired with a LSM710 confocal microscope (Zeiss) and stitched with Zen 2.1 (black, Zeiss). Confocal images were post processed with ImageJ (Version: 2.0). Cells were manually counted using the cell counter plugin (https://imagej.nih.gov/ij/plugins/cell-counter.html) for ImageJ.

### Statistical analysis

Statistical analysis was performed using Matlab (Mathworks) and Prism 8 (Graphpad). Unless otherwise indicated, normal distribution of the data was not assumed and non-parametric tests were performed. Values are presented as mean ± SEM unless stated otherwise. Box and whisker plots indicate median, interquartile range as well as the minimum to maximum value of the distribution. Statistical tests are mentioned in the text or figure legends. *, **, *** indicate P-values smaller than 0.05, 0.01 and 0.001, respectively.

---

**Figure S1:**
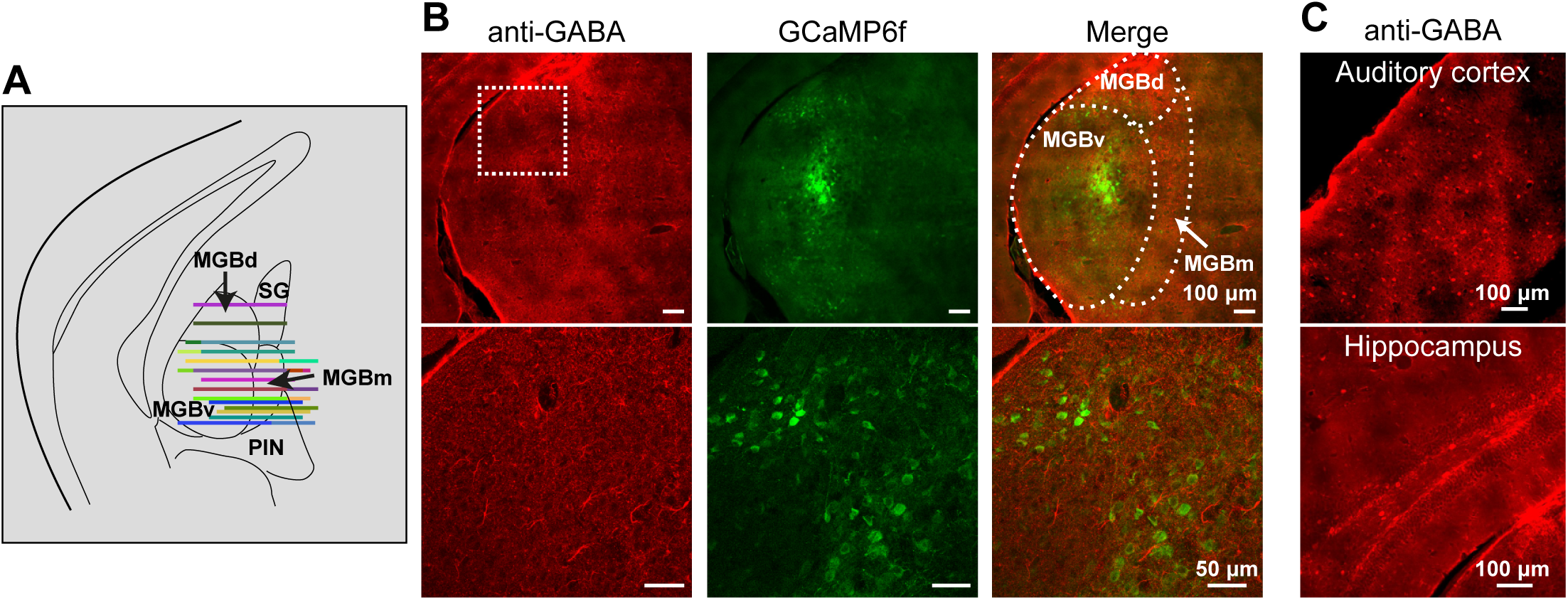
Imaging of excitatory cells in MGB. **A)** GRIN lens front (horizontal lines) of all mice (N = 21). **B)** Immunohistochemistry for GABA in MGB (left), GCaMP6f-expressing cells (middle) and merge (right). Bottom row: higher magnification of square indicated in top left image. GABAergic fibres are distributed widely in MGB while GABAergic somata are mainly absent (N = 2 mice). **C)** GABA-positive somata in auditory cortex and hippocampus using the same antibody as in B (N = 2 mice).

**Figure S2:**
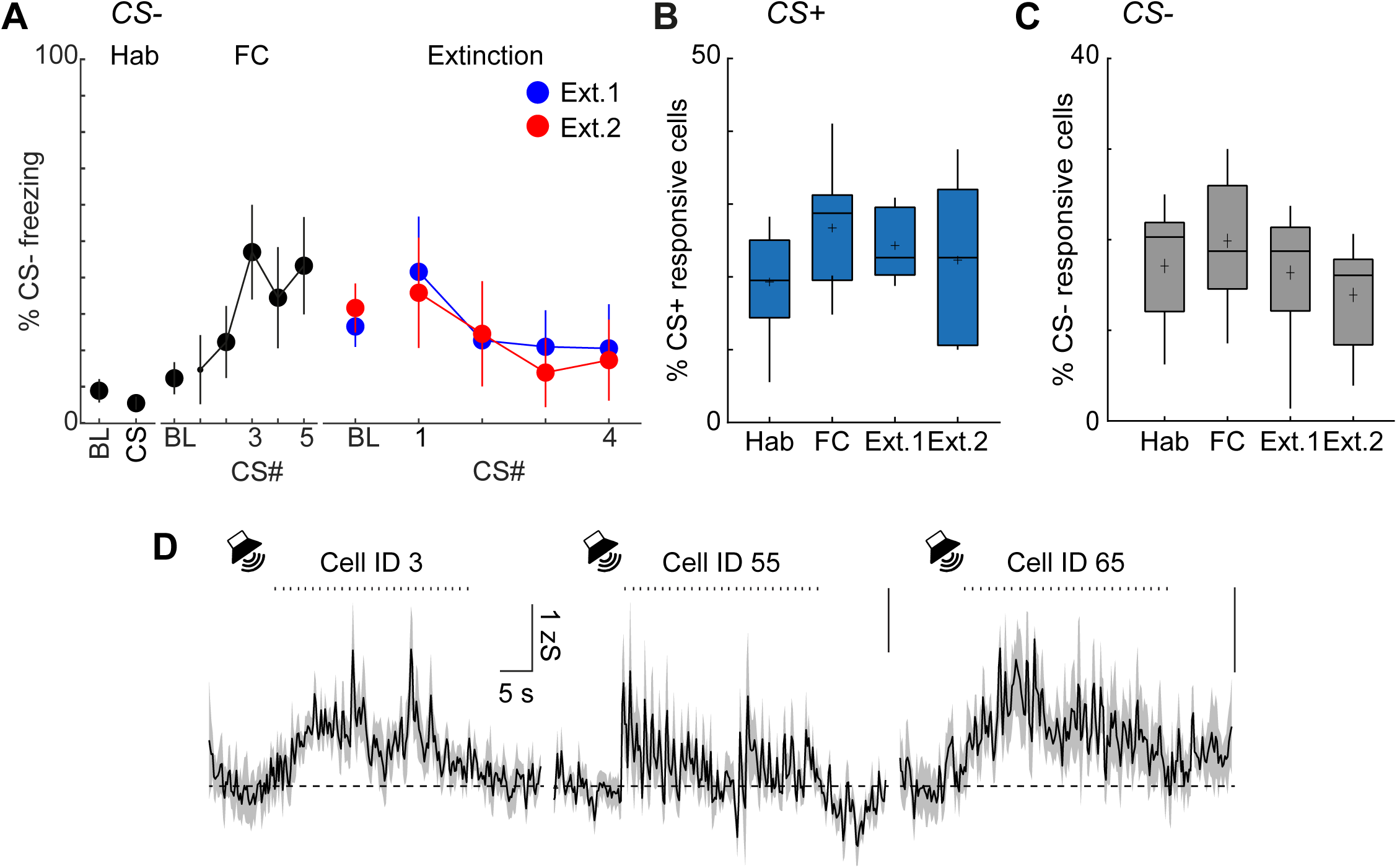
CS+ responsive cells, CS-freezing and CS-responsive cells. **A)** Freezing to the CS-during auditory fear conditioning (N = 15 mice). **B)** Percentage of CS+ responsive cells across all four imaging days (Friedman test, p > 0.05). **C)** Percentage of CS-responsive cells across all four imaging days. Hab = 17 ± 2%, FC = 20 ± 2%, Ext.1 = 16 ± 2%, Ext.2 = 14 ± 2%. Friedman test, p > 0.05. **D)** CS-Ca^2+^ responses of three example MGB neurons during habituation.

**Figure S3:**
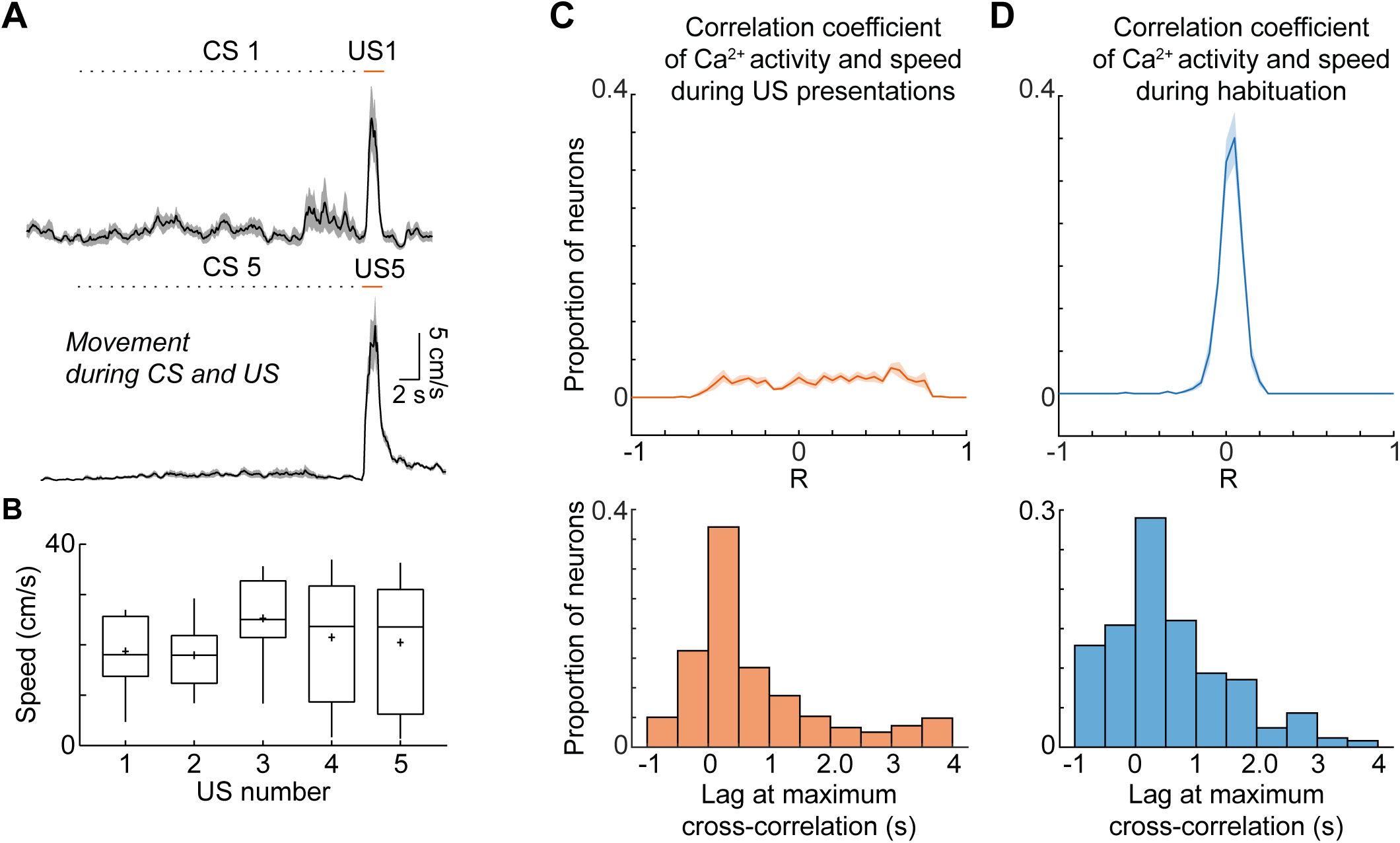
Correlation of neuronal activity and mouse movement. **A)** Average speed of mouse during first CS+-US presentation (top) and last CS+-US presentation (bottom). **B)** Average speed across the 5 US presentations (Kruskal-Wallis test, P > 0.05, N = 9 mice) **C)** Distribution of the maximum cross-correlation coefficients between Ca^2+^ activity of individual neurons and the mouse’s speed during the footshock US (n = 855 neurons from N = 9 mice, top). Lag of maximum cross-correlation coefficient (bottom). **D)** Distribution of the maximum cross-correlation coefficients between Ca^2+^ activity of individual neurons and the mouse’s speed during the habituation session outside of CS periods (n = 855 neurons from N = 9 mice, top). Lag of maximum cross-correlation coefficient (bottom).

**Figure S4:**
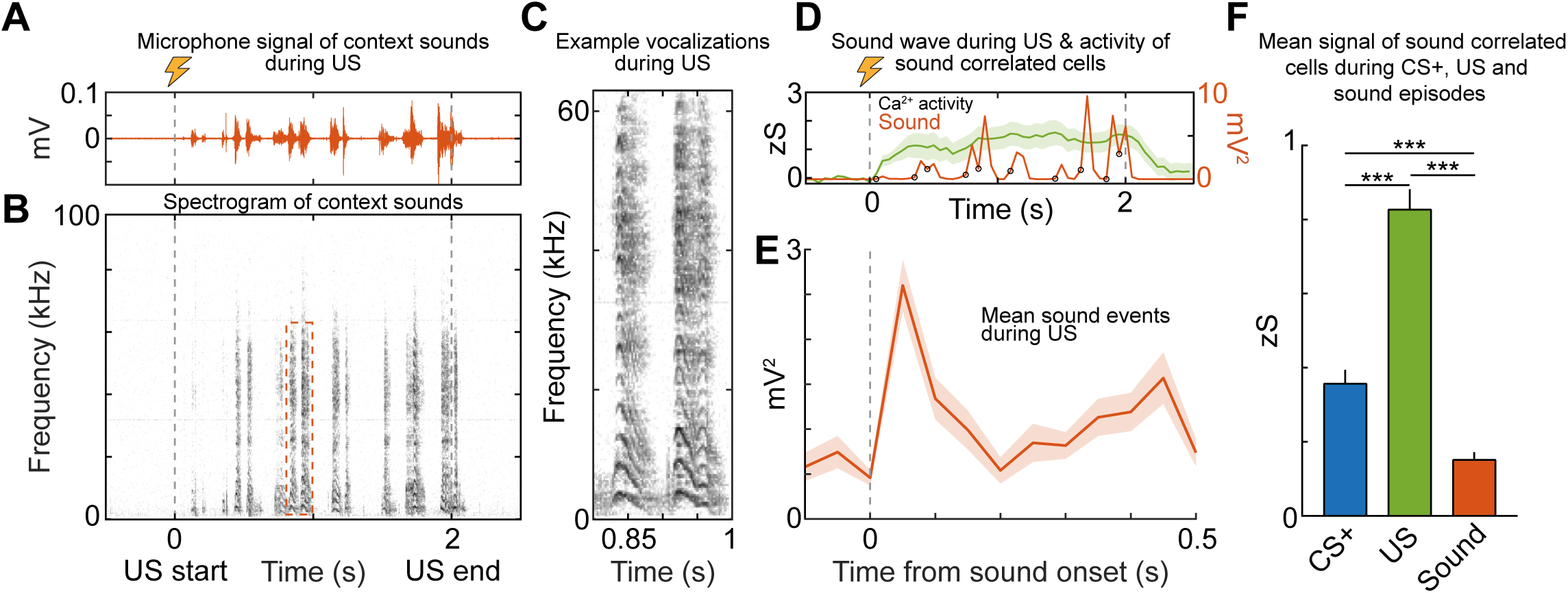
Ca^2+^ activity of MGB neurons is weakly correlated with self-evoked sounds during the US. **A)** Recorded sounds in the conditioning context during the two second US shock. **B)** Frequency spectrogram of the sound wave in A. **C)** Example low-frequency vocalizations during the US from the outlined box in B. **D)** Downsampled squared sound wave from A and the mean Ca^2+^ activity of cells which exhibited a cross-correlated Ca^2+^ response. Circles indicate the onset of sound events. **E)** Mean of onset-aligned detected sound events during all US presentations (n = 15 US presentations and 135 sound events from N = 3 mice, downsampled to match 20 Hz imaging frequency). **F)** Summary statistics of the mean Ca^2+^ response in Figure 2K (0-300 ms) indicate that US and CS responses are stronger than self-evoked sound responses (Kruskal-Wallis test with Dunn’s multiple comparisons test, all P < 0.001).

**Figure S5:**
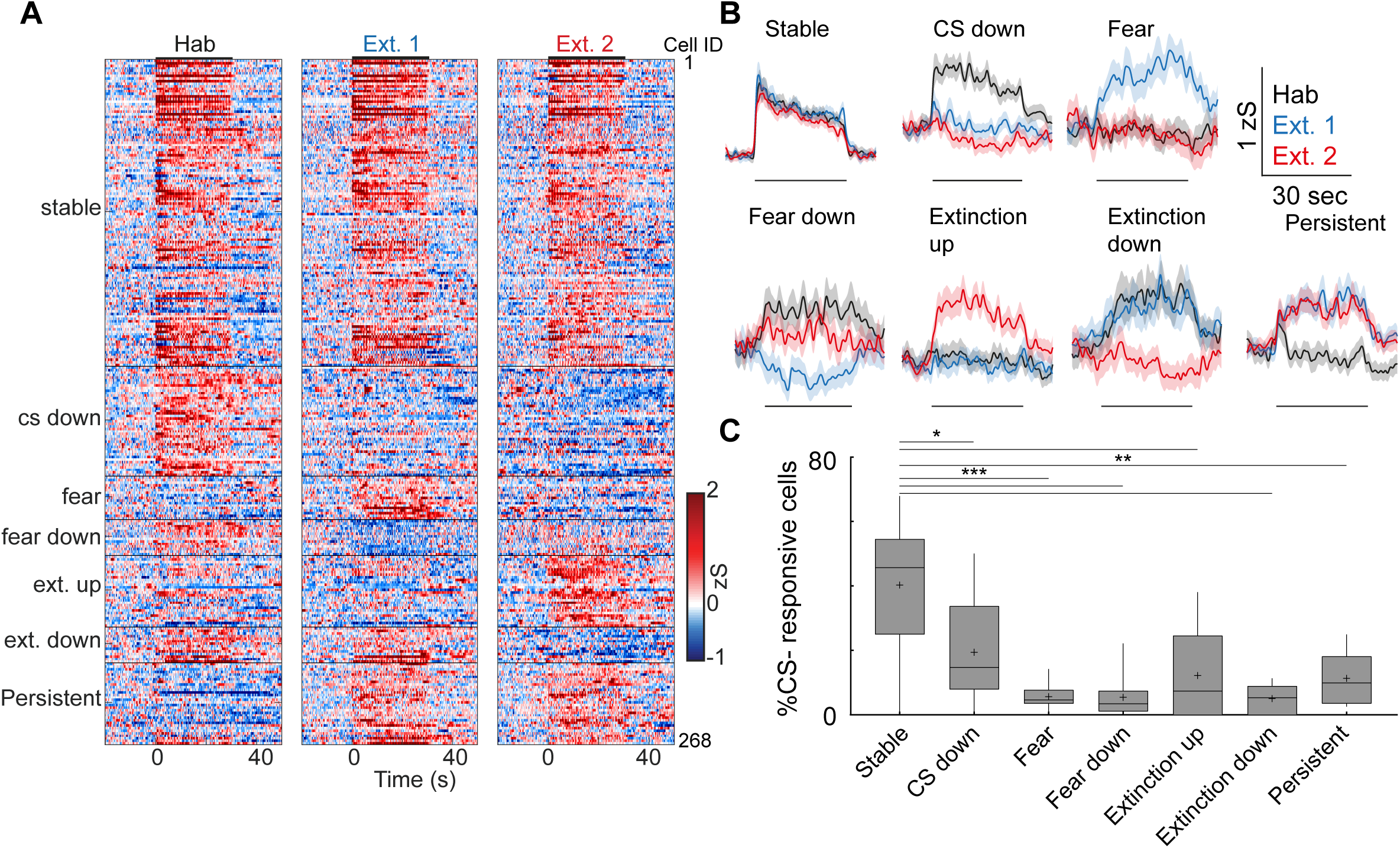
US and CS-response plasticity of individual MGB neurons. **A)** Response map of all CS-responsive neurons on the habituation, extinction 1 and extinction 2 days sorted by response type (n = 268 cells, N = 9 mice). **B)** Average Ca^2+^ traces of the different CS-response groups on the habituation (black), extinction 1 (blue) and extinction 2 (red) days. Traces represent mean ± s.e.m.. **C)** Quantification of CS-responses (Friedman test, p < 0.001, followed by Dunn’s multiple comparisons test, stable vs. extinction up p < 0.05; stable vs extinction down, p < 0.01; extinction down vs. persistent, p < 0.05).

**Figure S6:**
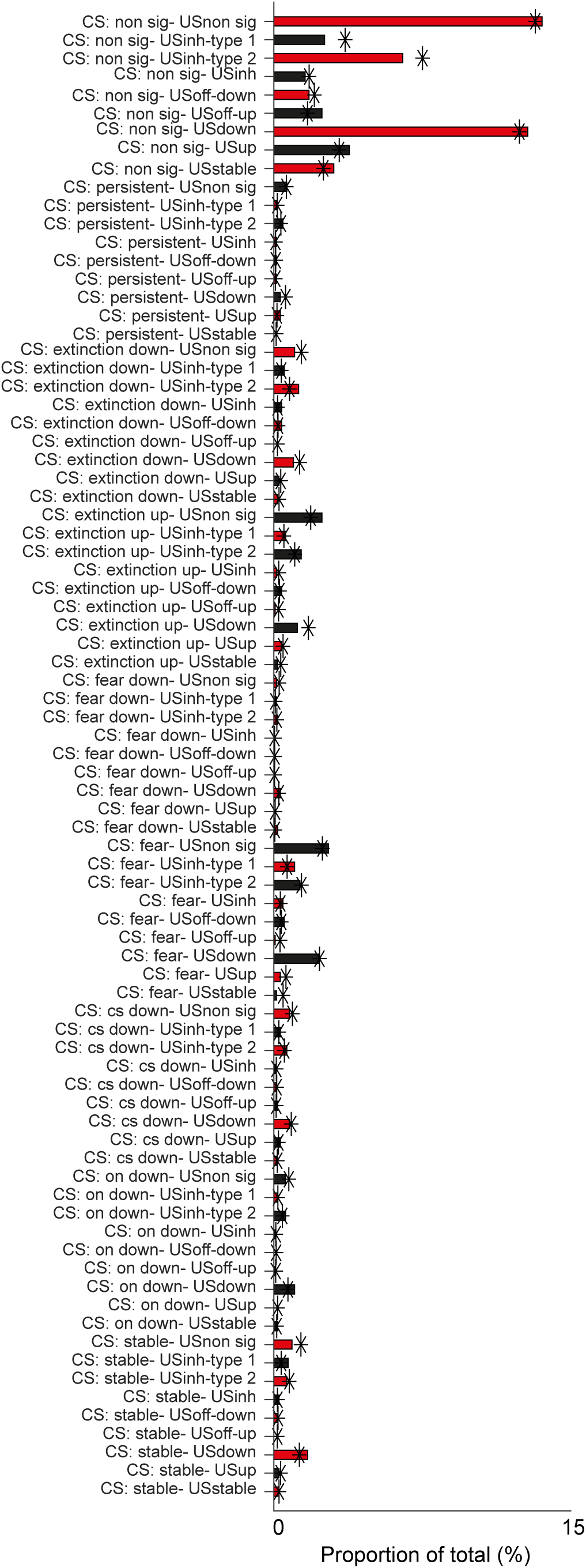
CS response type is not predictive of US plasticity. Proportion of overlapping subgroups of CS+ and US responsive cells (N = 9 mice). *indicates chance levels of finding overlapping groups based on the product of the proportions of individual subgroups.

**Figure S7:**
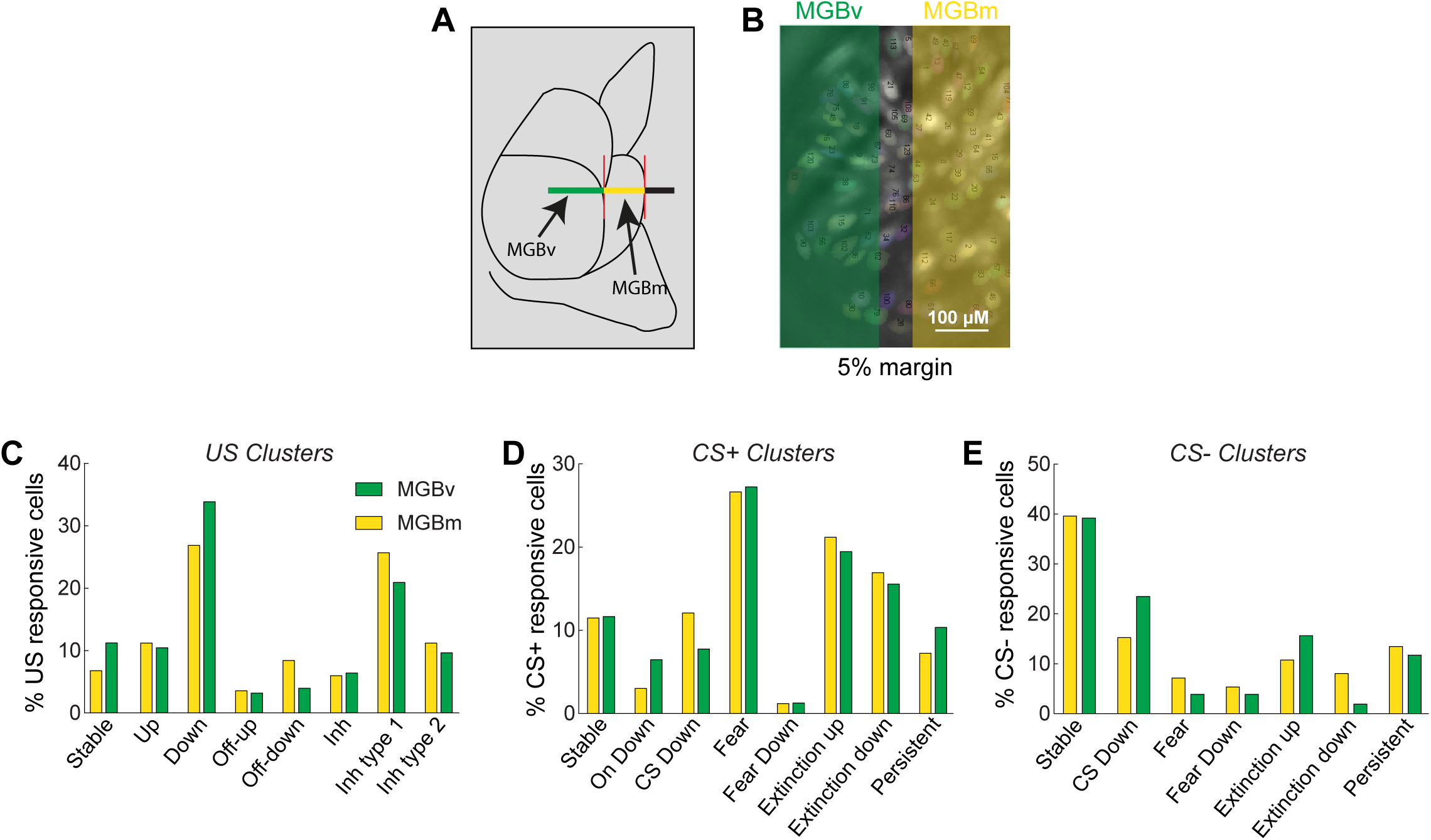
Similar CS and US plasticity types between MGB subdivisions. **A)** Schematic of GRIN lens location above different MGB subdivisions. **B)** Field of view of A divided into subregions. **C-E)** Proportion of cells belonging to the different US plasticity types (C), CS+ plasticity types (D) and CS-plasticity types (E) for each subdivision.

**Figure S8:**
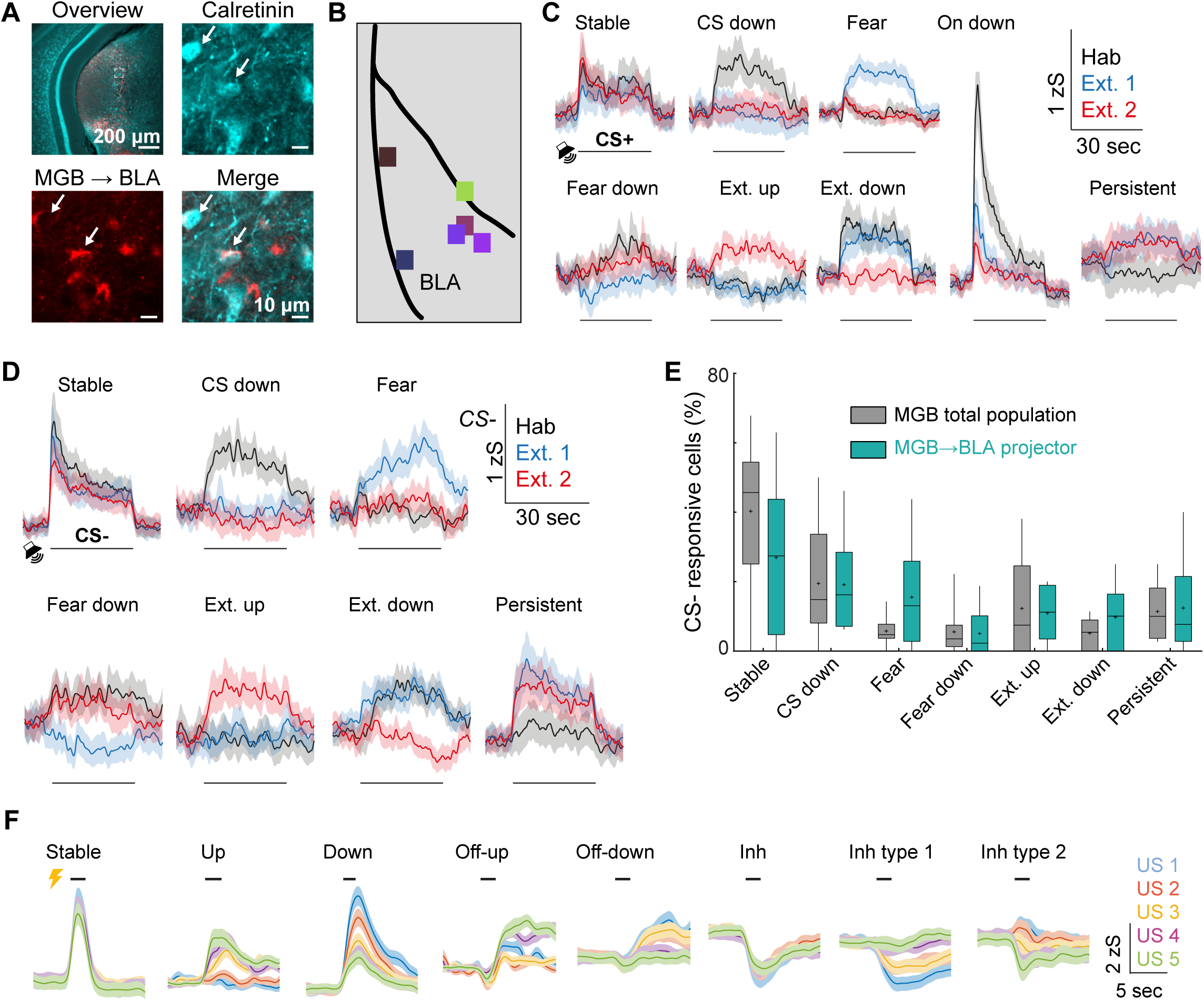
CS and US plasticity of MGB→BLA projection neurons. **A)** CTB labelling of MGB→BLA projection neurons and immunohistochemistry for calretinin. Arrows indicate double-positive cells. **B)** Center of virus injection sites of mice included in MGB→BLA projection neuron imaging (N = 6 mice). **C)** Average Ca^2+^ traces of CS+ plasticity subgroups of MGB→BLA projection neurons. Mean ± s.e.m.. Horizontal lines indicate CS+ period. **D)** Average Ca^2+^ traces of CS-plasticity subgroups of MGB→BLA projection neurons. Mean +/- s.e.m.. Horizontal lines indicate CS-period. **E)** Quantification of CS+ plasticity subgroups in the total MGB population (N = 9 mice) and MGB→BLA projection neurons (N = 5 mice). 2-way ANOVA, P > 0.05 **F)** Average Ca^2+^ traces of US plasticity subgroups of MGB→BLA projection neurons. Mean ± s.e.m.. Horizontal lines indicate US period.

**Figure S9:**
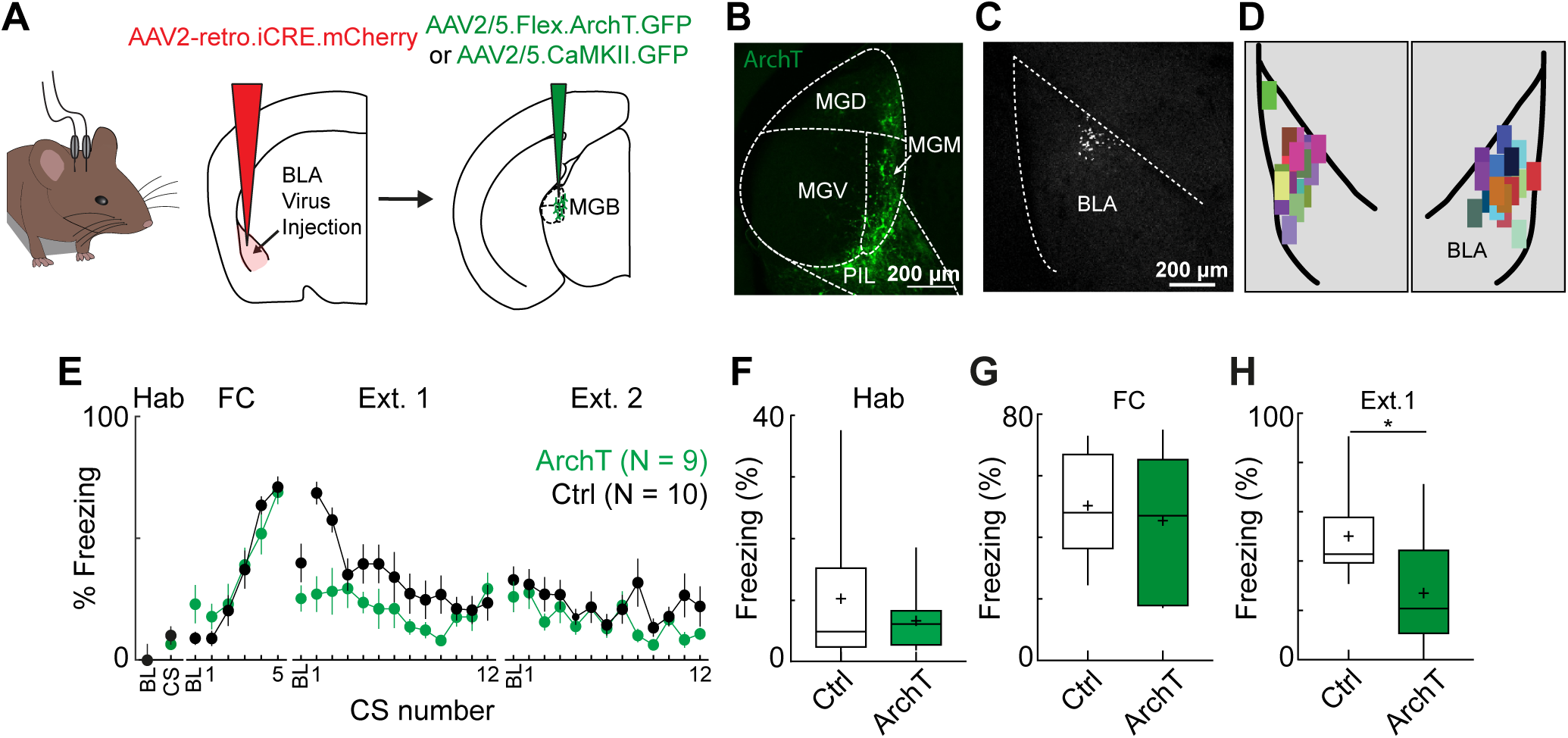
Optogenetic inhibition of MGB→BLA projection neurons reduces consolidation of fear learning. **A)** Viral expression strategy for optogenetic manipulation of MGB→BLA projecting neurons. **B)** Confocal image of ArchT-expressing MGB→BLA projection neurons. **C)** Example of latex beads in the basolateral amygdala (BLA). **D)** Center of rAAV2-retro.icre injection sites in BLA (N = 19 mice). **E)** Freezing across the fear conditioning paradigm for ArchT (green, N = 9 mice) and control mice (black, N = 10 mice). **F)** Quantification of freezing on the habituation day (Mann-Whitney test, p > 0.05). **G)** Average freezing of control and ArchT-expressing animals at the end of the fear conditioning paradigm (left, FC, freezing to last two CS, control: 50 +/- 5 % freezing, ArchT: 45+/- 8 % freezing, p > 0.05, Mann-Whitney Test). **H)** Average freezing of control and ArchT-expressing animals upon fear recall during early extinction 1 (Ext. 1, freezing during first four CS, control: 50 +/- 6 %, N = 10 mice, ArchT: 27 +/- 7 %, N = 9 mice, p < 0.05, Mann-Whitney test).

**Figure S10:**
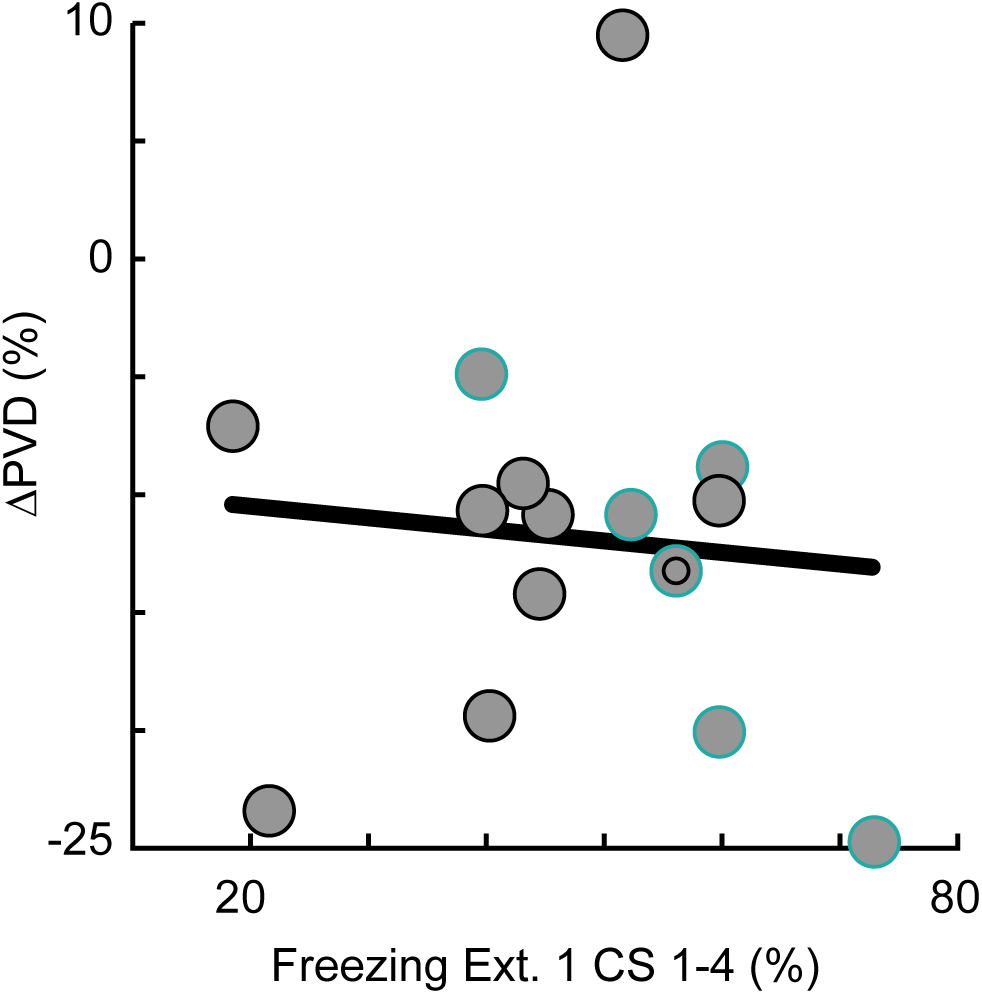
PVD change and fear learning are not correlated. Scatter plot of change in population vector distance (ΔPVD) and pre-extinction freezing behaviour for N = 15 mice. Black dots: Total MGB population. Cyan dots: MGB→BLA projector population. Line: Linear regression (R^2^ = −0.07, p > 0.05).

